# Sexually antagonistic environments and the stability of environmental sex determination

**DOI:** 10.64898/2026.03.08.710415

**Authors:** Erga Peter, Carl Veller, Pavitra Muralidhar

## Abstract

Just as sexually antagonistic genetic variants have different effects on male and female fitness, environmental conditions too can have sexually antagonistic fitness effects. Such ‘Charnov–Bull effects’ have been invoked to explain the origin and persistence of environmental sex determination (ESD), which allows development of each sex in the environments in which it has a comparative advantage. Here, we study different forms of Charnov–Bull effects to characterize how they shape the evolution and stability of ESD. We show that the precise functional form of Charnov–Bull effects can generate large differences in the vulnerability of ESD systems to the invasion of sex-biasing alleles, as well as in the fate of those alleles if they invade. For some configurations of Charnov–Bull effects, strong sex-biasing alleles are likely to spread to intermediate frequencies, rather than to fixation, resulting in ‘mixed’ ESD systems in which large genetic effects segregate. Overall, our results indicate that the precise nature of Charnov–Bull effects can play a crucial role in the evolutionary dynamics of ESD.

## 1 Introduction

Sexual antagonism occurs when males and females have different fitness optima for some phenotype but cannot evolve independently to these optima because they share much of their genome. This constraint represents a fundamental form of intragenomic conflict, and is common in species with separate sexes owing to differences between male and female morphology, physiology, and life histories [1–5].

As a common form of intragenomic conflict between the sexes, sexual antagonism is thought to play a critical role in the origin and stability of systems of genetic sex determination (GSD) [6–9]. Linkage between sexually antagonistic alleles and sex-determining mutations can drive the fixation of new sex-determining haplotypes and thus transitions to new sex-chromosome systems [6, 8–10]. Conversely, the presence of sexually antagonistic alleles on sex chromosomes can stabilize GSD systems by imposing fitness costs on the new sex-determining genotypes that necessarily arise during an evolutionary transition of the sex-determining system [6, 11]. Sexual antagonism is therefore a powerful evolutionary force that can drive both the origin and maintenance of GSD.

Although typically conceptualized in genetic terms—with males and females deriving different fitness outcomes from the same alleles—the logic underlying sexual antagonism readily extends to cases in which the sexes derive different fitness outcomes from the same environmental conditions. In many ectothermic species, for instance, the temperature at which an embryo develops influences its fitness differently if it develops as male or female [12–14]. Higher temperatures might accelerate growth, for example, which could increase the eventual mating success of males but have little effect on survival or reproduction of females. Such sex-specific impacts of environmental variables on fitness, which we refer to as as ‘Charnov–Bull’ effects [15], parallel the basic logic of sexual antagonism, differing only in that they operate through environmental rather than genetic variation.

Unsurprisingly, therefore, Charnov–Bull effects have been implicated as a principal mechanism underlying the evolution and maintenance of environmental sex determination (ESD). In ESD systems, an environmental cue, such as temperature during early development, determines whether an individual develops as male or female [6]. ESD is found across diverse animal clades, notably among reptiles and fish, and might have been the ancestral state for vertebrates [16–21]. By having sex determination depend on an environmental variable, ESD allows each sex to develop in those environments in which it has relatively higher fitness. In the example above, for instance, where warmer developmental temperatures improve fitness in males to a greater extent than in females, an ESD system that associates male development with high temperatures and female development with low temperatures maximizes fitness, compared to a GSD system in which sex determination would be random with respect to the developmental environment.

The key phenotype in ESD systems is the threshold: a value of the environmental variable below which an individual develops as one sex and above which they develop as the other sex [6, 14, 22]. Evidence across many ESD species indicates that thresholds are heritable quantitative traits with polygenic architectures [14, 23–28]. The evolutionarily stable threshold in an ESD system reflects the relationship between male and female fitness across environments, as well as the distribution of environments experienced by developing embryos [22,29,30]. The polygenicity of thresholds implies that they can adapt rapidly to changes in the environmental distribution, for example in response to climate change [13, 14, 26, 28].

Despite the intrinsic advantage of ESD in tailoring sex allocation to Charnov–Bull fitness effects, and the potential for ESD systems to adapt in response to environmental change, evolutionary transitions from ESD to GSD are much more common than the reverse, as reflected in the patchy phylogenetic distribution of ESD [17,20,21,31]. Several theories have been developed to explain how transitions from ESD to GSD might occur, for example via the spread of a major sex-determining allele linked to a sexually antagonistic allele [10], or the gradual strengthening of a sex-biasing haplotype composed of multiple linked threshold-affecting and sexually antagonistic alleles [32]. These theories implicitly assume that the genetic form of sexual antagonism outweighs its environmental form (i.e., Charnov–Bull effects), and they have led to a belief that ESD is inherently unstable, with transitions from ESD to GSD an inevitable consequence of genetic sexual antagonism.

However, recent comparative and genomic work suggests that transitions from ESD to GSD may be less straightforward than these theories imply [16, 33, 34]. Many apparent GSD systems show evidence of environmental influences and, conversely, many ESD systems show evidence of strong genetic effects [28,35–37]. Thus, while existing theories have sought to explain the conditions under which sex-determining alleles can invade, often with the implicit assumption that invasion of sex-determining alleles will be followed by their fixation and thus transitions to GSD [10, 32], the growing evidence for ‘mixed’ ESD–GSD systems suggests that these theories may not capture the full dynamics of transitions between ESD and GSD.

The stability of an ESD system will clearly depend to some degree on the nature of the Charnov– Bull effects that maintain it, as well as the distribution of environmental and genetic variation underlying the sex-determining system. Here, we present a general analysis of the invasion and the post-invasion dynamics of alleles that bias development towards one sex or the other in an ESD system. First, we derive conditions for the evolutionarily stable threshold values of an ESD system under a set of Charnov–Bull fitness functions. Given these, we then derive the conditions under which a sex-biasing, sexually antagonistic haplotype—i.e. a genetic sex determiner—can invade the ESD system. Finally, we characterize the dynamics that follow the initial invasion of such a haplotype, focusing on whether the haplotype spreads to fixation or to an intermediate frequency, and how the ‘background’ genetic value of the threshold it displaces adapts to accommodate its spread. Our interest is especially in the conditions under which the haplotype comes to be held in a long-term polymorphism, resulting in a mixed ESD–GSD system.

Our results reveal that the precise functional form of Charnov–Bull effects can have a strong influence on the dynamics at each stage in this process, and thus on the stability of ESD systems and the potential for mixed ESD–GSD systems.

## 2 Charnov–Bull effects

### 2.1 The source of Charnov–Bull effects

A wide range of environmental variables can differentially influence male and female fitness, and thus give rise to Charnov–Bull effects. Examples include temperature, salinity, light, pH, and population density [6, 18, 38, 39]. Among vertebrates, the environmental variable most commonly associated with sex determination is temperature [14, 17]. Temperature-dependent (or temperature-sensitive) sex determination (TSD) is common in fish and reptiles, and temperature-induced sex reversal has further been observed in several amphibian species previously thought to have exclusively GSD [14, 40, 41]. One possible mechanism for how temperature might induce Charnov–Bull effects is through a direct effect on early rates of development or growth, with different consequences for eventual male and female fitness [42]. For example, if higher temperatures accelerate growth and thus lead to larger eventual body size, and if female fecundity is particularly sensitive to body size, then females might benefit more than males from development in high temperatures. Alternatively, temperature could act indirectly, serving as a proxy for seasonal timing, with Charnov–Bull effects arising if there are sex-specific optima for hatching time during the nesting season [12, 43]. Recent experimental work supports the possibility that, in many vertebrate systems, temperature can act through a combination of the timing of maturation and sex-specific survival rates [40]. Thus, in general, the relationship between the environmental variable and Charnov–Bull effects can be mediated via diverse features of the life history of the species in question.

### 2.2 The shape of Charnov–Bull effects

Given the environmental variable underlying Charnov–Bull effects in a particular system, the shapes of the functions relating this environmental variable to male and female fitness will determine the parameters of the system of ESD that will evolve. In vertebrates, where temperature typically serves as the environmental cue, three general patterns of TSD have been observed, and are often assumed to reflect three distinct forms of Charnov–Bull effects [6, 14].

The first two patterns, found in many fish and reptiles, involve males developing at high temperatures and females at low temperatures, or vice versa (Fig. 1). These patterns are consistent with Charnov–Bull effects in which the male fitness function lies above the female fitness function for temperatures greater than some value, and below the female fitness function for temperatures smaller than this value, or vice versa. Such a ‘single crossing’ scenario might obtain, for example, if male fitness is more sensitive to temperature than female fitness is throughout the temperature range. The evolutionarily stable threshold in these systems is the temperature at which both sexes have equal fitness (refs. [29, 30] and below). In other words, at the evolutionarily stable threshold temperature, an individual should be indifferent between developing as male or as female.

**Figure 1.**
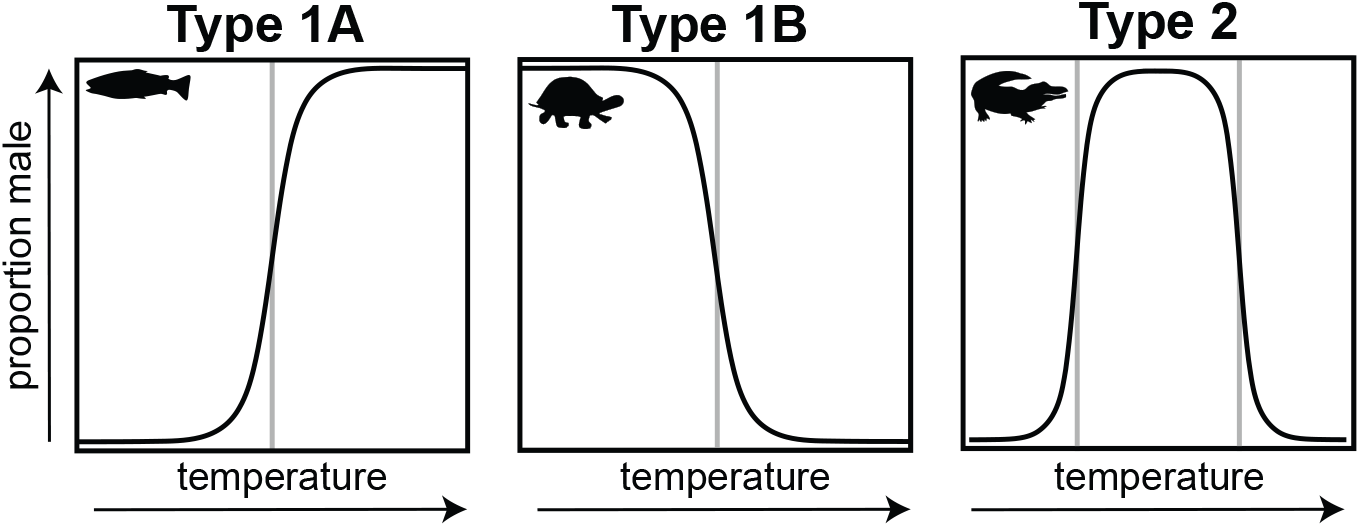
Patterns of temperature-dependent sex determination (TSD) in vertebrates.

In the third TSD pattern in vertebrates, males develop at intermediate temperatures and females develop at low and at high temperatures (Fig. 1). This pattern is consistent with Charnov–Bull effects in which male fitness lies above female fitness between two temperatures, and female fitness lies above male fitness below the lower and above the higher of these two temperatures. Such a ‘double crossing’ scenario might obtain if female fitness is a U-shaped function and male fitness an inverse-U-shaped function of temperature [12]. The evolutionarily stable thresholds in this system are the two temperatures at which the male and female fitness functions cross; at these temperatures, individuals are indifferent between developing as male or as female. Note that such two-threshold systems might appear to be single-threshold systems if the range of commonly observed temperatures does not overlap one of the thresholds.

Despite the central importance of Charnov–Bull effects in driving the evolution of ESD, little is known about their precise functional forms in nature. This is partly owing to the inherent difficulty of measuring fitness in natural populations (and here, across different developmental environments), and partly to the fact that, to estimate full Charnov–Bull functions, we must measure the fitness of each sex both in the environments in which it does ordinarily develop and in environments in which it does not. Exacerbating these difficulties is the fact that ESD is more common in long-lived species, the many overlapping generations in which can smooth out sex-ratio fluctuations caused by year-to-year environmental variation [44]. In such species, it is necessarily harder to measure lifetime fitness.

Despite these difficulties, several studies have successfully detected, and in some cases shone light on the functional form of, Charnov–Bull effects. Warner and Shine [12] combined artificially-induced sex reversal with long-term field experiments in the Jacky dragon, *Amphibolurus muricatus*, to quantify the impact of developmental temperature on lifetime reproductive success in males and females. They observed a distinct nonlinear pattern: male reproductive success was maximized at intermediate temperatures, whereas female reproductive success was maximized at both high and low temperatures. This Charnov–Bull pattern is consistent with the system of TSD found in this species (females high and low, males intermediate). Further work has indicated that the relationship between sex-specific reproductive success and temperature in this species is mediated, at least in part, by the association between incubation temperature and hatching time during the nesting season [12, 45, 46].

Spencer and Janzen [47] combined field measurements with experimental overwintering treatments to investigate Charnov–Bull effects in the painted turtle, *Chrysemys picta*. In this species, hatchlings spend winter in their natal nest, relying on maternal provisioning of yolk and other nutrients. Spencer and Janzen [47] collected eggs from wild nests and reared them under experimental treatments designed to mimic typical cold or warm overwintering conditions experienced in *C. picta* habitats. They found that male offspring more efficiently consume their nutritional reserves and respire in cooler temperatures, whereas females do so more efficiently in warmer temperatures. The ‘males low and females high’ system of TSD found in this species thus allows individuals to develop as the sex best suited to surviving winter given their initial nesting conditions.

Conover [43] studied TSD in the Atlantic silverside, *Menidia menidia*, using natural populations to investigate the effects of temperature on viability and reproductive success across the life cycle. He found that cooler temperatures correlate with earlier hatching times, earlier hatching correlates with larger body size at maturity, and body size more strongly correlates with gonad size (a proxy for reproductive success) in females than in males. While the relationship between temperature and fitness was not measured directly, the evidence suggests that female reproductive success is more sensitive to temperature than male reproductive success, with females having a comparative advantage in lower temperatures. The TSD system in this species, ‘females low and males high’, therefore allows individuals to use temperature as a cue of seasonality and develop as the sex whose reproductive success is relatively favored under those conditions.

As these examples illustrate, while the direction of Charnov–Bull effects can often be inferred (i.e., which sex benefits more from particular environmental conditions), their shape is much more difficult to establish. Nevertheless, the limited available evidence suggests that Charnov–Bull functions can be nonlinear and complex. This indicates that theoretical models of the evolution of ESD should consider a broad range of functional forms for Charnov–Bull effects.

## 3 Conditions for evolutionary stability of ESD thresholds

We are interested in the conditions under which a system of ESD can be invaded by sex-determining alleles, and the subsequent fate of these alleles, that is, whether they spread to fixation or to stable intermediate frequencies (resulting in a ‘mixed’ system). As a baseline, we need to characterize the properties of the system of ESD given (i) its set of Charnov–Bull fitness functions and (ii) the distribution of environments experienced by developing embryos. This involves determining the set of evolutionarily stable thresholds (which in practice will be just one or two) and the pattern of sex determination in each of the intervals that these thresholds delimit. The calculations in this section largely follow prior work [29, 30, 32], but they will be useful in the sections that follow, so we include them here for completeness.

We consider a variable *t* which is distributed across developmental environments within a breeding season according to the density *g*(*t*). For simplicity, we refer to this variable as incubation temperature, but it could be any environmental variable on which sex determination might be based. The viabilities—interpreted as probabilities of surviving to reproductive age—of a male and a female who develop at temperature *t* are *V*_*m*_(*t*) and *V*_*f*_ (*t*). The assumption that environmental effects on fitness are mediated through survival (rather than sexual competitiveness, say) is not crucial, but simplifies the analysis. We assume the functions *g, V*_*m*_, and *V*_*f*_ to be smooth. Each zygote is characterized by a set of heritable thresholds *θ* which delimit the temperature intervals in which they would develop as male and as female (in the simplest case, each zygote possesses a single threshold, developing as one sex at temperatures above the threshold and as the other sex at temperatures below the threshold).

Consider a set Θ of thresholds shared by every member of the population (along with a rule specifying which sex is produced in each inter-threshold interval). Label the set of male-determining temperatures ℳ and the set of female-determining temperatures ℱ. Then the numbers of males and females that survive to reproductive age are proportional to

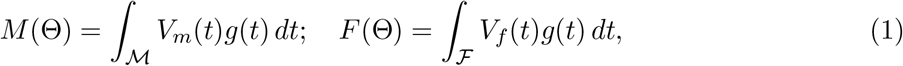

and so the expected reproductive success of a male and a female who develop at temperature *t* are proportional to

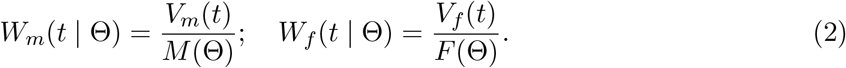

For a set of thresholds Θ^∗^ (and associated sex-determining rule) to be evolutionarily stable (ESS), each threshold *θ*^∗^ in the set must satisfy the condition

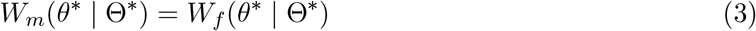

(refs. [6,22,29,30,32] and SI Section S1). That is, the expected reproductive success of the two sexes must be equal at each threshold. Because reproductive success incorporates both sex-specific viability and sex-ratio dependent reproductive value, the ESS thresholds generally do not correspond to points of intersection of the viability functions *V*_*m*_(*t*) and *V*_*f*_ (*t*) [6, 22] (Fig. 2).

**Figure 2.**
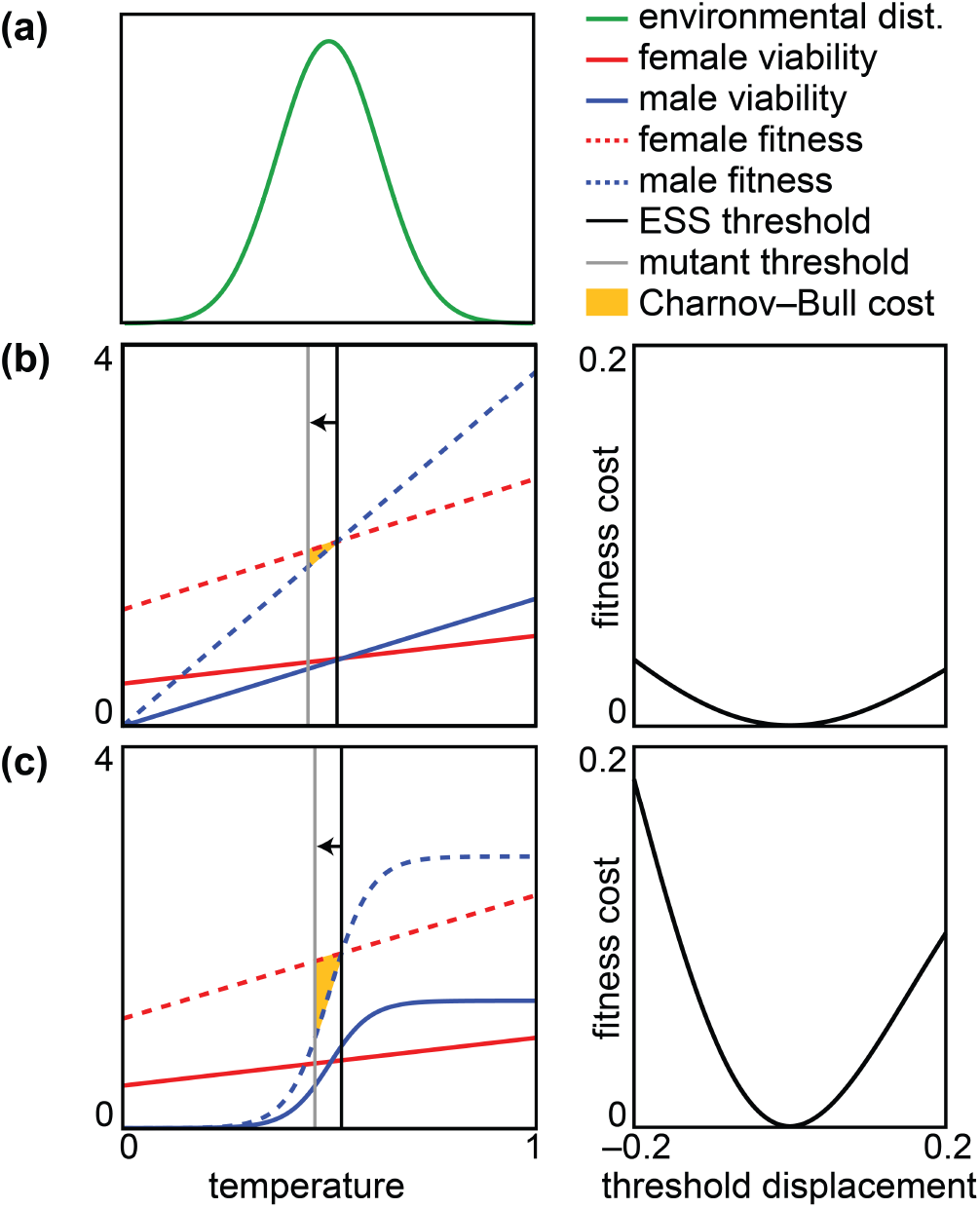
Fitness costs to alleles that displace thresholds from their evolutionarily stable values. (a) Here, temperature is assumed to follow a truncated normal distribution on the normalized range [0, 1]. (b) Male and female viability increase linearly with temperature, male viability more steeply so, supporting a ‘males high, females low’ ESD system. The threshold temperature determines the sex ratio after viability selection, which in turn determines the fitnesses of males and females at each temperature. The ESS threshold is such that male and female fitness, determined this way, cross at the threshold temperature. A mutant allele that displaces the threshold from its ESS value causes its bearers to develop as the lower-fitness sex in the interval between the displaced and ESS thresholds. The associated ‘Charnov–Bull’ fitness cost is the area of the yellow triangle between male and female fitness in this interval, weighted by the environmental distribution. The cost is larger for greater displacements of the threshold. (c) Male viability increases logistically with temperature, while female viability is the same linear function as in b. The resulting ESS threshold and primary sex ratio are similar to those in b. However, the nonlinear male fitness function leads to larger costs for alleles that displace the threshold from its ESS value, relative to the linear scenario in b. Parameters can be found in SI Section S6.

The second condition for evolutionary stability concerns the sex-determining rule associated with Θ^∗^: for each threshold *θ*^∗^ ∈ Θ^∗^, if 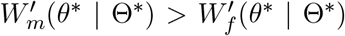, then females are produced in the temperature interval below *θ* and males are produced in the interval above *θ*, while if 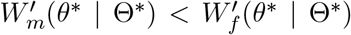, then males are produced in the interval below *θ*^∗^ and females are produced in the interval above *θ* (SI Section S1).

Apart from some pathological cases (where the fitness functions ‘kiss’ at certain temperatures), these conditions, together with our assumption that *V*_*m*_, *V*_*f*_, and *g* are smooth, guarantee regular alternation of male and female production across adjacent intervals.

### Single-threshold systems

A sufficient condition for stability of a single-threshold system is that male and female viability are monotonic in temperature, and the viability of one sex is always more sensitive to temperature than the viability of the other sex. Thus, for example, if 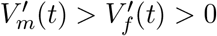 across the full range of experienced temperatures *t*, then there is a unique threshold value *θ*^∗^ such that the TSD system in which males develop at temperatures above *θ*^∗^ and females develop at temperatures below *θ*^∗^ is stable. In such cases, the primary sex ratio is skewed towards the sex that develops in the lower-viability environments (females in this example) [30, 48].

### Two-threshold systems

In contrast, for more complicated, non-monotonic fitness functions, multiple thresholds can exist in a stable TSD system. Here we focus on the empirically motivated case in which female fitness is U-shaped and male fitness is inverse-U-shaped, as observed in the Jacky dragon, *A. muricatus* [12]. In this case, males are favored over an intermediate range of environments, and so the stable TSD system has two thresholds, 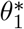 and 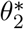. The ESS condition (3) must hold at each of these threshold temperatures:

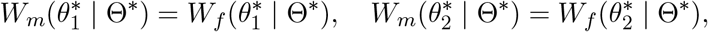

and in this case, 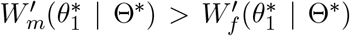 while 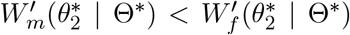. In such nonmonotonic cases, the intuition about primary sex ratios derived from monotonic single-threshold systems does not necessarily apply: depending on the precise fitness functions and the distribution of temperatures, either sex may be overrepresented at equilibrium. Consequently, when Charnov– Bull fitness relationships are non-monotonic, unbiased primary sex ratios are insufficient to exclude that sex is environmentally determined [49].

Having established the ESS structure of ESD systems under Charnov–Bull fitness functions, we now consider whether these systems can be invaded by sex-determining alleles.

## 4 Fitness cost to a threshold-biasing allele in an ESD system

In the section above, we derived conditions for what might be called the ‘internal’ stability of an ESD system—that is, stability with respect to perturbations of the system that involve parameters endogenous to it (i.e., thresholds and sex-determining rules). We now turn to the ‘external’ stability of ESD systems under Charnov–Bull fitness effects, specifically their stability with respect to the potential invasion of alleles with exogenous fitness effects that push the system towards one of GSD. We are particularly interested in alleles (or haplotypes) that simultaneously predispose their bearers to develop as one sex or the other, by effecting a change in their thresholds, and exogenously improve the fitness of the sex to which they bias the development of their bearers. We begin by characterizing the fitness costs that such alleles suffer by displacing their bearers’ thresholds from the ESS values.

In general, threshold-biasing alleles are genetic variants that alter the thresholds of their carriers. For example, in a single-threshold, ‘males high, females low’ TSD system, an allele that shifts its carriers’ thresholds downwards by a small amount behaves as a small-effect male-determining allele: its carriers are slightly more likely than average to develop as male, but still develop as female in sufficiently cool environments. Were the allele to have a larger effect on the threshold, causing its carriers to develop as male across the full range of commonly experienced temperatures, it would function as a large-effect male-determining allele. If such an allele were to invade and spread to high frequency, all else equal, the sex-determining system would effectively transition from ESD to GSD.

Two forces constrain the spread of threshold-biasing alleles: (1) sex-ratio costs and (2) Charnov– Bull costs. Sex-ratio costs arise because a threshold-biasing allele will, all else equal, skew the population sex ratio toward the sex overproduced by that allele, reducing its own reproductive value and impeding its spread. However, these costs will be negligible when the allele is rare, since then it has only a negligible effect on the population sex ratio. Sex-ratio costs can therefore be ignored when considering potential invasion of the allele, but not when considering its subsequent spread to higher frequency—although as it spreads, the sex ratio bias caused by the allele might be mitigated by adaptive response of the ‘background’ threshold phenotype (we consider such dynamics in Section 5.3 below).

Charnov–Bull costs to threshold-biasing alleles arise because, by shifting thresholds from their ESS values, these alleles cause their bearers to develop as one sex in certain environments where the other sex has higher fitness (by the ESS condition (3) above). These costs pertain even when the allele is rare, and therefore impede its invasion. To quantify the Charnov–Bull fitness cost to a rare allele that displaces a threshold from its ESS value, we assume that all individuals in the population initially have thresholds equal to their ESS values (the set Θ^∗^).

Consider a rare allele that has the additive effect of decreasing one of the thresholds by an amount *δ*, from its ESS value *θ*^∗^ to *θ*^∗^ − *δ* in heterozygotes and to *θ*^∗^ − 2*δ* in homozygotes. We assume, if the system is a multi-threshold one, that the displaced threshold *θ* − 2*δ* does not overshoot any of the thresholds that lie below *θ*^∗^ in the set of ESS thresholds Θ^∗^. Suppose, without loss of generality, that 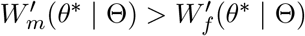 in the resident system, so that males have higher fitness than females (and are therefore produced) in the interval above *θ*, while females have higher fitness than males (and are therefore produced) in the interval below *θ*^∗^. Individuals who carry the rare mutant allele (almost always as heterozygotes) develop as female in the interval below *θ*^∗^ − *δ* and as male in the interval above *θ*^∗^ − *δ*; the allele therefore reduces their expected fitness because it causes them to develop as male at temperatures between *θ*^∗^ −*δ* and *θ*^∗^, where females have higher fitness. The associated fitness cost is

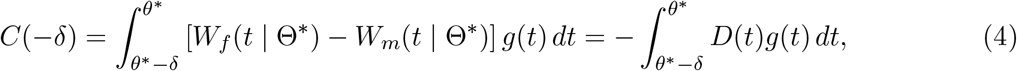

where *D*(*t*) = *W*_*m*_(*t* | Θ^∗^) − *W*_*f*_ (*t* | Θ^∗^) is the difference between male and female fitness at temperature *t* in the resident system, and is negative in the interval (*θ*^∗^ − *δ, θ*^∗^). This fitness cost corresponds to the area of the triangle formed by the female and male fitness functions between *θ*^∗^ − *δ* and *θ*^∗^, weighted by the density of temperatures *g*(*t*) in this interval (Fig. 2) [32].

If the threshold displacement *δ* is small, we show in SI Section S2 that Eq. (4) simplifies to

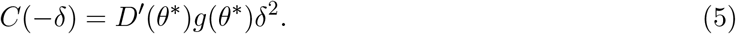

Note that 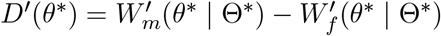 is positive, by assumption.

The calculations above are for a threshold-decreasing allele, but are easily modified for the case of a threshold-increasing allele (which in this system would predispose its bearers to develop as female). In general, the cost to an allele that increases a threshold from its ESS value *θ*^∗^ to *θ*^∗^ + *δ*, assuming this increased threshold not to overshoot any thresholds above *θ*^∗^ in the system, is

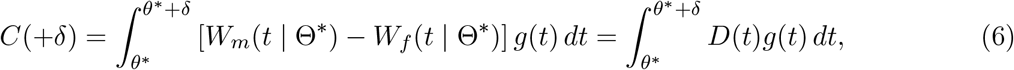

which for small-effect alleles can again be approximated by

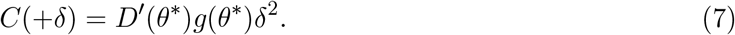

(If instead *D*^′^(*θ*^∗^) *<* 0, so that females are produced in the interval above *θ*^∗^ and males in the interval below, then identical cost functions to those above would be found by defining 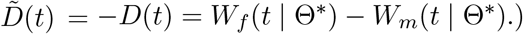.

The fitness costs to alleles that displace thresholds from their ESS values under Charnov–Bull fitness functions are seen to grow quadratically with the size of their effects on the threshold, consistent with the interpretation of these costs as areas of triangles formed between the male and female fitness functions (Fig. 2). Thus, alleles with very small threshold-biasing effects are, all else equal, very weakly selected against, and might segregate at high frequency. This perhaps helps to explain the substantial genetic variation that is often observed for thresholds, despite stabilizing selection on their values [14, 23, 24].

For small-effect alleles, it is the difference in the slopes of the male and female fitness functions at the relevant threshold, *D*^′^(*θ*), that governs the fitness cost to the allele. This is a consequence of the fact that, in the close vicinity of the threshold, these functions can be approximated as linear, and so their slopes alone govern the area of the triangle they form between the displaced and ESS thresholds (Fig. 2). This linear approximation breaks down for large-effect alleles that displace the threshold far from its ESS value, and the Charnov–Bull fitness costs they suffer come to depend on more details of functional form of the male and female fitness functions in the interval [*θ*^∗^, *θ*^∗^ + *δ*), and in particular on their curvature.

### Examples

To illustrate how different Charnov–Bull functions can influence the invasion of large-effect threshold-biasing alleles, we consider two representative forms of these functions, both leading to single-threshold ESD systems in which males develop at high temperatures and females at low. The first is a simple linear case, which serves as a baseline for comparison and has been studied previously [30, 32]. In this scenario, both male and female viability increase linearly with temperature, but male viability does so with a steeper slope (Fig. 2b). In the second scenario, male and female viabilities again increase monotonically with temperature: the female viability function is the same as in the first scenario, but male viability follows a logistic curve (Fig. 2c). This second scenario is inspired by the relationship between size and fitness in the Atlantic silverside, *M. menidia* [43]. Temperature is assumed to follow a truncated normal distribution across the range [0, 1] (Fig. 2a), and the male viability functions have the same minimum and maximum values across this range in the two scenarios. The single-threshold, ‘males high, females low’ ESD systems that evolve under these two Charnov–Bull scenarios have nearly identical ESS thresholds and thus primary sex ratios (Fig. 2).

Despite these similarities in the observed characteristics of the two systems, the fitness costs to threshold-biasing alleles can differ substantially between them. In the linear case, Charnov–Bull costs increase gradually with the size of the threshold displacement, but in the nonlinear case, these costs accelerate once an allele’s displacement of the threshold exceeds a certain magnitude (Fig. 2).

We also consider a two-threshold system, inspired by the Charnov–Bull effects estimated for the Jacky dragon, *A. muricatus* [12], with male fitness at intermediate temperatures and female fitness maximized at both temperature extremes, leading to a ‘females high and low, males intermediate’ system of ESD (Fig. 1). Under the conservative assumption that threshold-affecting alleles displace only one of the two thresholds, the curvature of both the male and female fitness functions in this scenario causes Charnov–Bull costs to rise sharply with the magnitude of the threshold displacement (Fig. 4). For alleles that displace both thresholds at once, these costs would be even greater. Thus, although the precise scaling of the fitness costs to threshold-biasing alleles depends on details of the fitness functions, ESD systems based on this form of Charnov–Bull effects may be particularly resistant to the invasion of large-effect threshold-biasing alleles.

## 5 Invasion and spread of a sexually antagonistic, threshold-biasing haplotype in an ESD system

The previous section established that threshold-biasing alleles suffer fitness costs owing to Charnov– Bull effects, and are therefore unlikely to invade and spread in isolation unless they have very small effects on the threshold. However, these alleles may also be associated with exogenous fitness advantages that outweigh their costs, and can invade on that basis. Of particular interest here are sexually antagonistic genetic effects—where an allele increases the fitness of one sex but reduces the fitness of the other sex—which have been proposed as a mechanism by which threshold-biasing alleles can invade an ESD system [10, 32].

We consider an additive threshold-biasing allele that decreases one of the thresholds in Θ^∗^ by an amount *δ*, from its ESS value *θ*^∗^ to *θ*^∗^ − *δ* in heterozygotes and to *θ*^∗^ − 2*δ* in homozygotes. We further assume that males are produced in the interval above *θ* in the resident system—i.e., that *D*^′^(*θ*^∗^) *>* 0—so that the allele is male-biasing. Our calculations are easily adapted for female-biasing alleles as well. We assume that the threshold-biasing allele is associated with sexually antagonistic fitness effects, either through pleiotropy or through linkage with sexually antagonistic alleles at other loci. If the latter, for simplicity, we assume complete linkage between the threshold-biasing and sexually antagonistic alleles, so that they form a threshold-biasing, sexually antagonistic haplotype. We call this haplotype *H*, and the resident haplotype *h*. The sexually antagonistic allele has selection coefficient *s*_*m*_ in males and *s*_*f*_ in females, and its fitness effects are multiplicative, so that, relative to *hh* males, *Hh* and *HH* males have their viabilities multiplied by 1 + *s*_*m*_ and (1 + *s*_*m*_)^2^, while *Hh* and *HH* females have their viabilities multiplied by 1 + *s*_*f*_ and (1 + *s*_*f*_)^2^, relative to resident *hh* females.

### 5.1 Conditions for invasion of the haplotype

For the sexually antagonistic, threshold-biasing haplotype *H* to invade, its direct effects on fitness in heterozygotes, via *s*_*m*_ and *s*_*f*_, must be net positive and must outweigh the fitness cost the haplotype suffers owing to Charnov–Bull effects, *C*(−*δ*), calculated above. The general invasion condition is

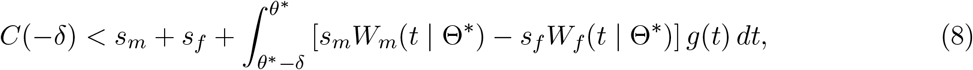

as shown in ref. [32] and SI Section S3. We further show in SI Section S3 that, when the threshold shift *δ* is small, and the direct fitness effects of the haplotype, *s*_*m*_ and *s*_*f*_, are also small (specifically O(*δ*^2^); i.e., on the order of the Charnov–Bull fitness cost of the haplotype), condition (8) simplifies to

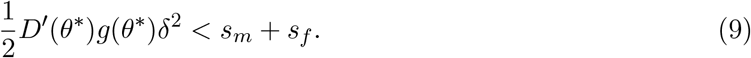

For sexually antagonistic effects, where *s*_*m*_ and *s*_*f*_ differ in sign, Eq. (9) reveals that, if the male-biasing threshold effect of the haplotype *H* is small, then it can invade whether it is male beneficial (*s*_*m*_ *>* 0 *> s*_*f*_) or female beneficial (*s*_*f*_ *>* 0 *> s*_*m*_). On the other hand, if the male-biasing threshold effect of *H* is large, Eq. (8) shows that *H* is much more likely to invade if it is male-beneficial; indeed, since *W*_*m*_(*t* | Θ^∗^) *< W*_*f*_ (*t* | Θ^∗^) in the interval (*θ*^∗^ − *δ, θ*^∗^), for the integral on the right-hand side of Eq. (8) to be positive requires *s*_*m*_ *>* 0 *> s*_*f*_ . This is sensible: if the haplotype ensures that it is much more likely to be found in males, then it will more readily invade if it specifically increases male fitness.

The calculations above can easily be modified for a threshold-increasing haplotype, which predisposes its bearers to develop as female. We show in SI Section S3 that the general invasion condition for a female-biasing haplotype with threshold effect +*δ* and selection coefficients *s*_*m*_ and *s*_*f*_ is

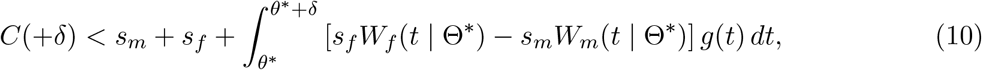

which, for small *δ*, can again be approximated by Eq. (9).

The calculations are also easily adapted to the case where *D*^′^(*θ*^∗^) *<* 0, so that males are produced in the interval below *θ*^∗^ and females above, by simply swapping the labels ‘*m*’ and ‘*f*’ in the calculations above, and defining 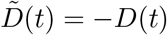.

### 5.2 Conditions for long-term polymorphism of the haplotype

If the haplotype invades, it will spread either to an intermediate frequency or to fixation. If it spreads to an intermediate frequency, a mixed ESD–GSD system will result. Here, we derive the conditions for long-term polymorphism of the haplotype.

The logic that we apply is straightforward. Suppose the haplotype *H* satisfies the invasion criterion (8), so that it will invade a resident *hh* system. We next consider a situation where the population is fixed for *H* and, given this, the ESD system has adjusted to its new ESS. If *h*, the haplotype in the original system of ESD, can invade this new system in which *H* is fixed, then we conclude that *H* would not have been able to spread to fixation in the first place, and instead will come to be held in a long-term polymorphism with *h*.

To determine whether *h* can invade a system in which *H* is fixed, we must characterize the properties of this system. In particular, we need to know what the ESS thresholds are conditional on *H* being fixed. Fortunately, these are just the same ESS thresholds Θ^∗^ from the original system in which *h* was fixed. That is, for each threshold *θ*^∗^ ∈ Θ^∗^, the ‘background’ genetic contribution to this threshold has increased by an amount 2*δ* in adaptive response to the threshold-decreasing effect −2*δ* of the fixed haplotype *H* (assuming that there is sufficient genetic variation in the background threshold to effect this change). The reason is that, with *H* fixed, all males have their viabilities multiplied by the same factor (1 + *s*_*m*_)^2^, and all females have their viabilities multiplied by the same factor (1 + *s*_*f*_)^2^. If the thresholds of the system were Θ^∗^, then the numbers of males and females that survive to reproductive age (Eq. 1) would be multiplied by these same factors, and so the fitnesses of a male and female at temperature *t* would be unchanged, relative to the original system in which *h* was fixed (Eq. 2). Since these fitness functions would be unchanged, and since Θ^∗^ is the ESS set of thresholds in the original system in which *h* was fixed, it is also the ESS set of thresholds in the new system in which *H* is fixed.

Therefore, the invasion conditions (8) and (10) from the system fixed for *h* apply to the system fixed for *H* as well. Relative to the sexually antagonistic, threshold-decreasing haplotype *H*, the original haplotype *h* can itself be thought of as sexually antagonistic and threshold-increasing, with an additive effect +*δ* on the threshold and with direct, multiplicative fitness effects 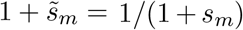 and 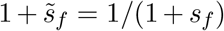 in males and females respectively. From Eq. (10), the condition for reverse invasion of *h* in a system fixed for *H* is therefore

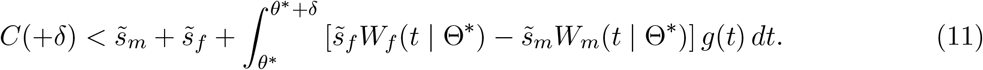

If the parameters of the system and the haplotype *H* are such that conditions (8) and (11) are satisfied, then we expect *H* to invade and spread to intermediate frequency, where it will subsequently be held in a long-term polymorphism with *h* (Figs. 3, S3).

**Figure 3.**
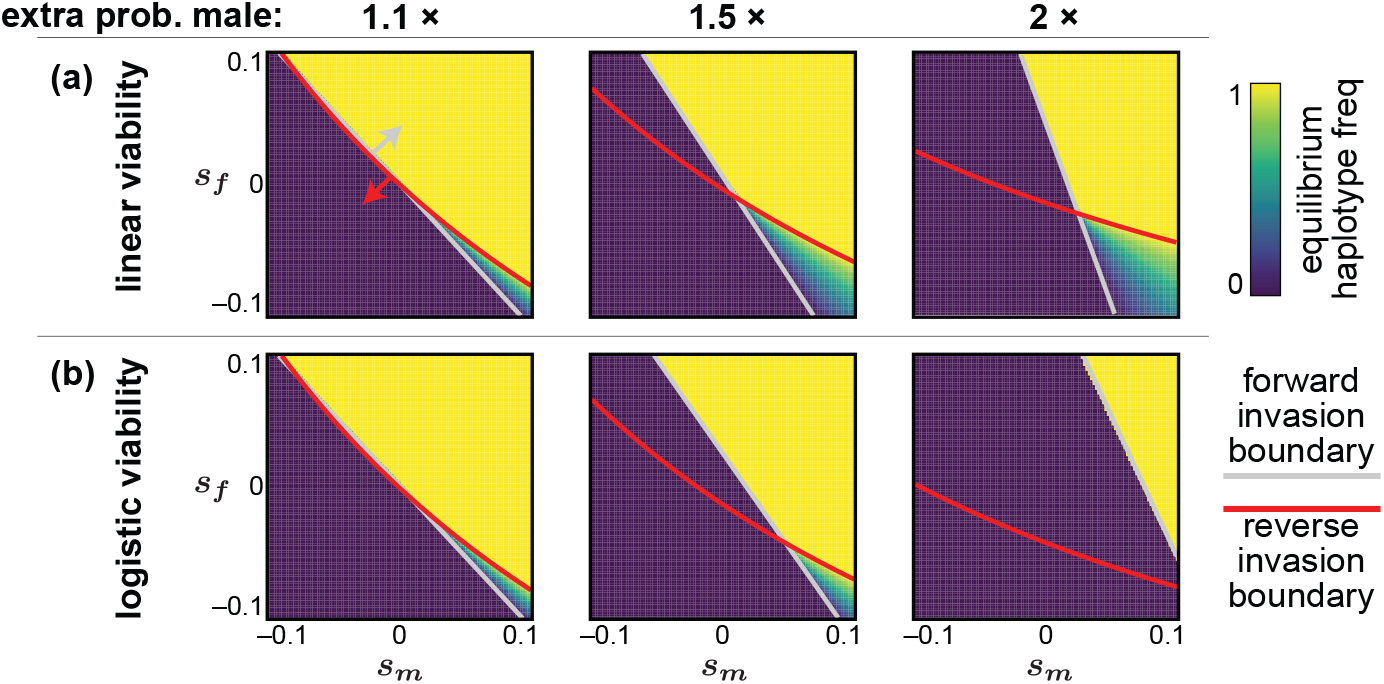
Conditions for invasion and polymorphism, and equilibrium frequencies, of sexually antagonistic, threshold-biasing haplotypes. The system in (a) is the same as that in Fig. 2b, with linear male and female viability functions, while the system in (b) is as in Fig. 2c, with a logistic male viability function. The mutant haplotype is assumed to be threshold-decreasing, and therefore to increase the probability that its bearers develop as male; columns correspond to the fold-change in the probability of developing as male, relative to individuals in the original ESS system. In parameter regions above the grey line, the mutant haplotype can invade a population fixed for the original haplotype. In regions below the red line, the original haplotype can reinvade a population fixed for the mutant haplotype. Regions above the grey and below the red line correspond to long-term polymorphism of the haplotype. Polymorphism is seen to be possible only in the sexually antagonistic region of parameter space, *s*_*m*_ *>* 0 *> s*_*f*_ . Within this region, in the linear Charnov–Bull scenario (a), large-effect haplotypes more readily invade than small-effect haplotypes, and are also more likely to spread to intermediate frequencies, resulting in ‘mixed’ ESD–GSD systems. In contrast, in the nonlinear Charnov–Bull scenario, haplotype invasion is less likely than in the linear case, and, moreover, large-effect haplotypes are less likely to invade and attain stable polymorphisms than small-effect haplotypes. For a zoomed-in view of the sexually antagonistic region of parameter space (*s*_*m*_ *>* 0 *> s*_*f*_), see Fig. S3.

If *δ, s*_*m*_, and *s*_*f*_ are small, Eq. (11) can be approximated by

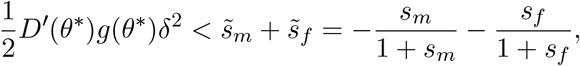

so that the condition for long-term polymorphism of the haplotype is

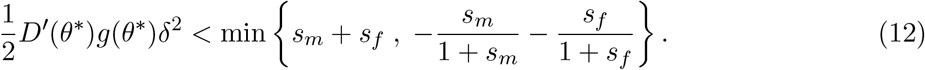

It follows from Eq. (12) that, for the haplotype eventually to be held at intermediate frequency, both terms on the right-hand side must be positive, and so the direct fitness effects of the haplotype *s*_*m*_ and *s*_*f*_ must be sexually antagonistic—that is, we must have that *s*_*m*_ *>* 0 *> s*_*f*_ or *s*_*f*_ *>* 0 *> s*_*m*_. Moreover, condition (12) is more likely to be met for larger-magnitude effects *s*_*m*_ and *s*_*f*_ (Fig. S3), mirroring classic results for long-term polymorphism of sexually antagonistic alleles [50–52].

Fig. 3 shows the parameter regions supporting long-term polymorphism of the haplotype *H* in the two monotonic, single-threshold ESD systems discussed earlier, which differ only in that the male viability function is linear in one scenario and nonlinear in the other. The regions supporting polymorphism of haplotypes with small effects on the threshold are broadly similar across the two systems, despite the difference in the shapes of the male viability functions. In contrast, the fate of larger-effect alleles differs substantially across the two systems, with the strong Charnov–Bull costs in the nonlinear system severely restricting the parameter space supporting polymorphism.

The same phenomenon occurs, with even more force, in the non-monotonic system we have studied. Conservatively assuming the haplotype to affect only one of the two thresholds, we see that the parameter space supporting eventual polymorphism rapidly shrinks as the threshold effect gets larger (Fig. 4).

**Figure 4.**
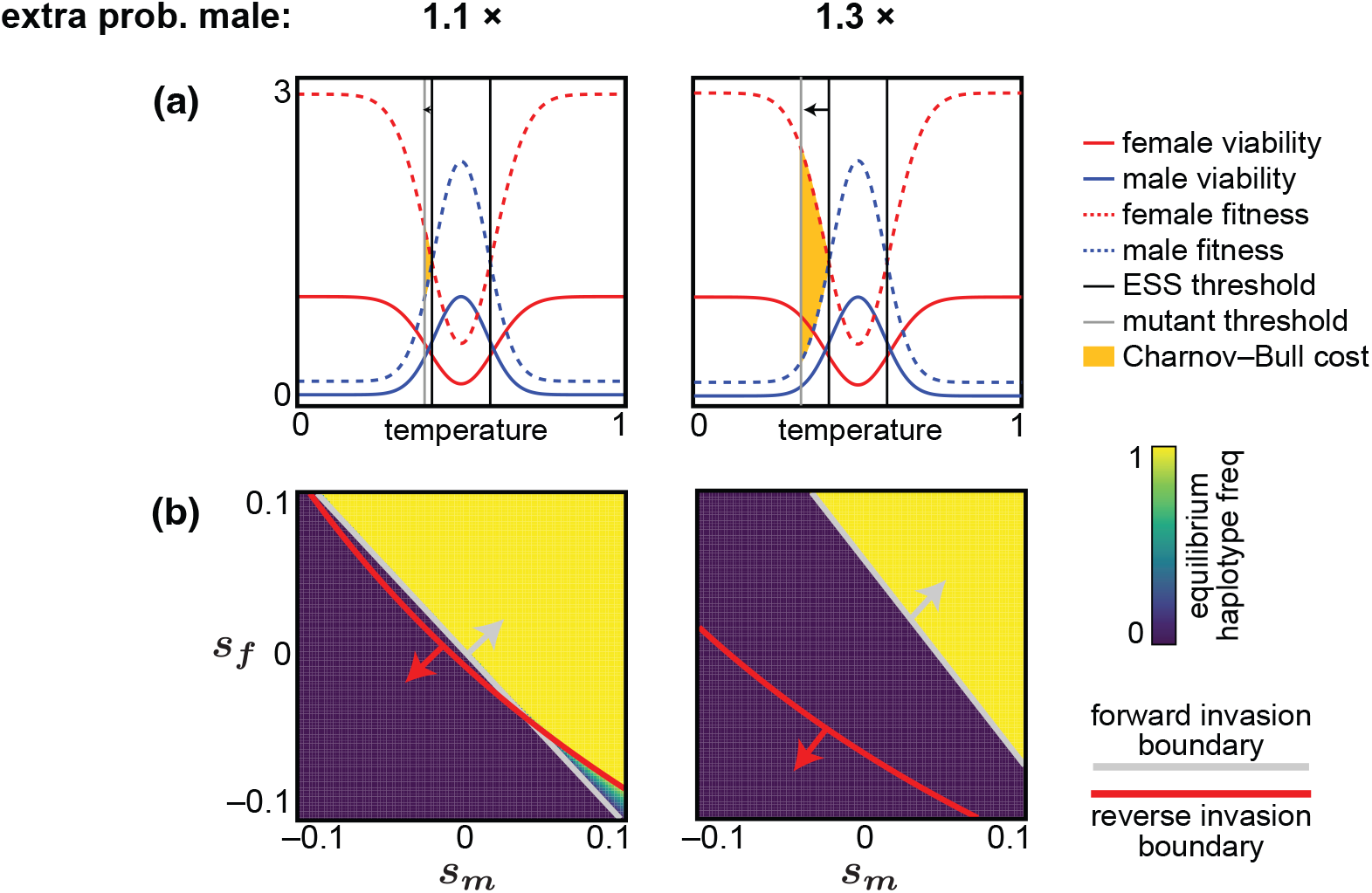
Invasion and polymorphism of threshold-biasing haplotypes in a two-threshold ESD system. (a) Charnov–Bull viability functions, and corresponding fitness functions, underlying a two-threshold ESD system in which males are produced at intermediate temperatures and females are produced at extreme temperatures. Mutant alleles that displace either threshold incur a fitness cost, as they produce the lower-fitness sex in the interval between the ESS and mutant threshold. This cost, represented by the area of the triangle between the male and female fitness functions bounded by the mutant and ESS thresholds, is especially severe in this case relative to that in Fig. 2, owing to the opposite-sign slopes of the male and female fitness functions at the ESS threshold (i.e., the triangle is wedge shaped rather than a sliver). (b) Owing to these large fitness costs associated with displacement of the threshold, a sexually antagonistic, male-biasing haplotype can invade and spread to intermediate frequency only in a small region of parameter space. Moreover, for large threshold effects (here, 1.3-fold increase in the probability of developing as male, vs. 1.1-fold), only those sexually antagonistic haplotypes with strong male benefits can invade, and the region supporting polymorphism collapses.

### 5.3 Equilibrium frequency of the haplotype

If conditions (8) and (11) hold, the sexually antagonistic, threshold-biasing haplotype *H* will invade and spread to intermediate frequency. Here, we characterize the frequency to which it will spread.

All else equal, the threshold-decreasing haplotype’s spread would cause the sex ratio to become more male biased. This generates selection for compensatory adaptation of the ‘background’ value of the threshold, via changes in allele frequencies at the many loci assumed to underlie genetic variation in the threshold in the original ESD system. The rate at which adaptation of the background threshold occurs in response to the haplotype’s spread will in turn affect the haplotype’s dynamics— for instance, if there were no response of the background threshold, then the sex ratio would become progressively more male biased, reducing the reproductive value of the haplotype and thus checking its spread.

To numerically characterize the equilibrium frequency of the haplotype, we simulate the dynamics assuming a constant, positive genetic variance for the background threshold. This assumption is consistent with a highly polygenic architecture, as thresholds have often been inferred to possess in natural systems [14, 23–28].

We begin with the haplotype *H* at a low starting frequency *ε*, with the genotypes *HH, Hh*, and *hh* in males and females in the Hardy–Weinberg proportions 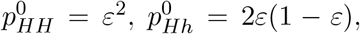, and *p*_*hh*_ = (1 − *ε*)^2^. The mean value of the background threshold starts at 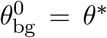, the ESS threshold value in the original system. From this starting configuration, we iterate the dynamics as follows. In generation *k*, the genotype frequencies among male and female zygotes are 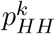, 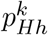, and 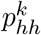. The mean value of the background threshold is 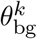, so that the mean thresholds among *HH, Hh*, and *hh* individuals are *θ*_bg_ − 2*δ, θ*_bg_ − *δ*, and *θ*_bg_ respectively. From these values, we calculate: (i) the frequencies 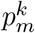 and 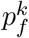 of the haplotype among males and females after selection and, from these, the frequencies of the three genotypes in male and female zygotes next generation, 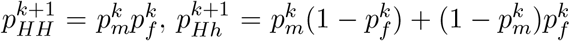, and 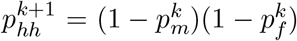; (ii) the selection gradient *S* acting on the background threshold, evaluated at its current mean value 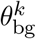, from which we determine the mean value of the background threshold next generation using the breeders’ equation, 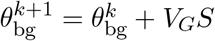. Details of these calculations can be found in SI Section S5. The process is iterated until convergence.

Fig. 3 shows the resulting long-term frequencies of the haplotype across the two monotonic, single-threshold ESD systems we have analyzed. The regions in which the haplotype attains intermediate frequency in our simulations are seen to fall between the boundaries defined by the conditions (8) and (11) calculated earlier. As with the regions supporting polymorphism, the equilibrium frequencies of haplotypes with small effects on the threshold are broadly similar across the two systems, but the equilibrium frequencies of haplotypes with large effects differ substantially.

Previous models have suggested that, if a threshold-biasing, sexually antagonistic allele invades and attains a stable polymorphic frequency, it could subsequently evolve to become both more strongly sex-biased and more sexually antagonistic, which could permit it to increase further in frequency [32]. Our results indicate that such post-invasion escalation is not guaranteed. As the magnitude of the threshold bias increases, the parameter space supporting polymorphism can contract rather than expand (as in Figs. 3b and 4). In such cases, the accumulation of stronger threshold-biasing effects on the haplotype will more likely lead to loss or fixation of the haplotype, rather than to stable polymorphism at higher frequency. Similarly, intensifying the sexually antagonistic effects of the haplotype may drive its fixation, rather than establishing it at a higher equilibrium frequency.

### 5.4 Impact of the environmental distribution

In all of our calculations above, the distribution of environments *g*(*t*) plays a central role. The prospects for invasion and spread of a sexually antagonistic, sex-biasing haplotype will therefore depend on the nature of this distribution, in addition to the nature of the Charnov–Bull effects. For example, the cost to a haplotype that displaces a threshold from its optimal value *θ*^∗^ to *θ*^∗^ *δ* is given by the area of the triangle formed by the female and male fitness functions between *θ*^∗^ *δ* and *θ*^∗^, but with each temperature *t* in this interval weighted by its density *g*(*t*) (Figs. 2, 4). Thus, if the density *g* places large weight on the temperature range [*θ*^∗^ − *δ, θ*^∗^) over which the haplotype produces the lower-fitness sex, the fitness cost to the haplotype will be correspondingly large, and it will be less likely to invade (Fig. S2). Conversely, if *g* places little weight on this range, the fitness cost to the haplotype will be small, because it ‘misassigns’ sex only in rarely experienced temperatures, and so the haplotype will be more likely to invade.

Therefore, just as the conditions for invasion and polymorphism of large-effect haplotypes can be sensitive to the precise shape of Charnov–Bull fitness effects, and particularly nonlinearity of these effects, so it can be sensitive to the precise shape of the environmental distribution *g*(*t*). Such sensitivity may contribute to intraspecific variation in sex-determining systems [34–36, 53], with differences in environmental distributions across a population’s range promoting stability of ESD in some parts but facilitating transitions to GSD in others. Since climate change is expected to alter the distributions of environments experienced by ESD species, accounting for these distributions— and geographic variation in their shape—will be important for understanding the persistence of ESD systems in the face of climate change [13, 54].

## 6 Discussion

Environmental sex determination has often been regarded as an evolutionarily unstable system, which will inevitably be replaced by genetic sex determination over time [31,32]. Our results suggest that this view may implicitly reflect assumptions about the simplicity of the shape of Charnov–Bull fitness relationships underlying ESD, with more complex Charnov–Bull relationships sometimes having a substantial effect on the the stability of ESD systems. In particular, we find that, while the potential for polymorphism of genetic sex-determining alleles with minor effects (i.e., weak threshold-biasing alleles) is relatively insensitive to the shape of the Charnov–Bull relationship, the potential for polymorphism of major-effect sex determining alleles (i.e., strong threshold-biasing alleles) can be constrained by nonlinearity of Charnov–Bull effects.

Given the impact they can have on the ultimate stability of an ESD system, our results indicate a need for accurate measurement of Charnov–Bull effects in natural systems. Progress in this area will require not only documenting sex ratios and/or developmental outcomes, but also quantifying how reproductive success varies with the developmental environment for each sex, and understanding the mechanistic basis of these relationships. Recent experimental and comparative studies in natural populations provide promising examples of how such work might proceed [36, 40, 46, 47, 55].

In considering the invasion and post-invasion dynamics of a sexually antagonistic, threshold-biasing haplotype, we assumed complete linkage between the sexually antagonistic and threshold-biasing alleles. Prior theoretical work has shown that, for a sexually antagonistic, threshold-biasing haplotype to overcome its Charnov–Bull fitness cost and invade an ESD system, tight linkage between its constitutive alleles is required [32]. Thus, our results can be thought of as providing an upper bound for the ease with which genetic sex determiners can invade ESD systems and spread.

Additionally, we have found that, when a two-threshold system is underlain by U-shaped fitness functions of opposite orientation in the two sexes (as has been estimated in a model ESD system, the Jacky dragon [12]), there is only a narrow region of parameter space in which a threshold-biasing sexually antagonistic haplotype can invade and spread to intermediate frequency. This is because the costs to such a haplotype from displacing a threshold from its ESS value are especially severe given this shape of fitness functions. As we have noted, the cost to a threshold-displacing allele can be visualized as the area of the triangle formed by the male and female fitness functions between the displaced and ESS thresholds, and in this case, the triangle is wedge shaped (Fig. 4) rather than a sliver, as it is in the case of monotonic fitness functions (Fig. 2). This observation suggests that two-threshold systems, when underlain by Charnov–Bull effects like those estimated in the Jacky dragon [12], might be more resistant to invasion of genetic sex determiners than single-threshold systems are. This might explain why two-threshold ESD is widespread in reptile clades [14, 21, 31].

An open question concerns the molecular and developmental basis of the threshold phenotype in ESD systems. In our analysis, we made conservative assumptions about how threshold-biasing alleles act—for example, assuming that such alleles shift only one boundary in two-threshold systems rather than shifting both boundaries simultaneously (in the same or opposite directions) [21,34]. Existing work suggests that genetic variation can influence thermal sensitivity and sex-determination pathways in diverse ways, but it is still unclear how different classes of mutations are most likely to affect the threshold phenotype in a given species [39, 56, 57]. Recent mechanistic models of temperature-dependent sex determination (e.g., [58, 59]) have proposed developmental frameworks for how temperature and genotype interact to produce sex, and offer a promising foundation for ultimately linking proximate mechanisms of sex determination with the population-genetic approaches to understanding Charnov–Bull relationships taken here.

Recent empirical data indicate that in some populations, genetic sex-determining variants with moderate effects may be segregating in systems previously thought to be purely ESD [37]. Such variants can generate patterns in which sex determination appears environmentally determined under some conditions but genetically determined under others. Characterizing these mixed ESD– GSD systems offers an important opportunity to study partial transitions between ESD and GSD, as well as the co-evolutionary dynamics between sex-biasing alleles and the threshold phenotype. Similarly, the increasing availability of large-scale population genomic data from ESD species provides a promising avenue for detecting and tracking the joint evolution of genetic and environmental contributions to sex determination [28, 60, 61].

We have focused on the evolution of the threshold phenotype as the primary adaptive response to invasion of sex-biasing alleles. However, in many species with ESD, nesting behavior—which can include nest-site choice, incubation timing, or maternal provisioning—can play a key role in influencing offspring sex ratios. To the extent that such nesting behaviors are heritable, they can act as an alternative or complementary adaptive mechanism by which ESD systems can buffer against the sex ratio distortion induced by the invasion and spread of sex-biasing alleles [46, 62, 63].

In conclusion, our work shows that the fate of an ESD system can be highly sensitive to the functional form of the Charnov–Bull effects that underlie it. Characterizing these effects empirically is therefore crucial for understanding the resilience of ESD systems to invasion of genetic sex determiners. This will be particularly important in the context of ongoing climate change, which will likely alter the distribution of environments experienced by populations and the relationship between those environments and sex-specific fitness.

## Acknowledgements

This work was supported by a Rosalind Franklin Young Investigator Award from the Gruber Foundation and the Genetics Society of America, awarded to PM.

## Supplementary Information for

**Sexually antagonistic environments and the stability of environmental sex determination**

Erga Peter, Carl Veller, Pavitra Muralidhar

## S1 ESS thresholds under Charnov–Bull fitness effects

Some quantitative environmental variable *t*, which for simplicity we will refer to as incubation temperature, is distributed according to the probability density *g*(*t*). Males and females incubated at temperature *t* have viability *V*_*m*_(*t*) and *V*_*f*_ (*t*) respectively. The functions *g, V*_*m*_, and *V*_*f*_ are assumed to be smooth. We assume that the population has non-overlapping generations and is large enough that stochastic effects can be ignored, that zygotes are assigned randomly to environments, and that surviving males and females mate randomly.

A system of environmental sex determination is defined by a set of temperature thresholds Θ = {*θ*_1_, *θ*_2_, …, *θ*_*n*_}, with *θ*_1_ *< θ*_2_ *<* … *< θ*_*n*_, and a corresponding rule for which sex is produced in each of the intervals delimited by these thresholds. Given *g, V*_*m*_, and *V*_*f*_, we are interested in determining the conditions that an ESD system must satisfy to be ‘internally’ evolutionarily stable, by which we mean stable with respect to invasion of other ESD systems (but not necessarily with respect to other ‘external’ forces, such as the appearance of sexually antagonistic alleles). To derive the conditions for internal stability of Θ (and the corresponding sex-determining rule), we largely follow the methodology of refs. [29, 30, 32].

Let ℱ and ℳ be the sets of environments in which females and males are produced, respectively, under the system. ℱ and ℳ are each a union of intervals delimited by the thresholds in Θ, and together constitute a partition of the temperature range. The sex ratio at the time of sex differentiation is

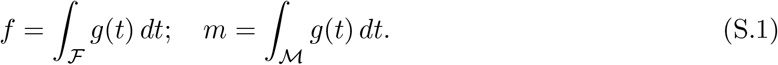

After viability selection, the numbers of males and females are proportional to

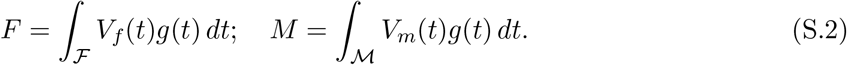

The average reproductive success of a surviving female is proportional to 1*/F*, while the average reproductive success of a surviving male is proportional to 1*/M* . Therefore, the expected fitnesses of a female and a male who develop at temperature *t*, taking into account viability and sexual selection, are

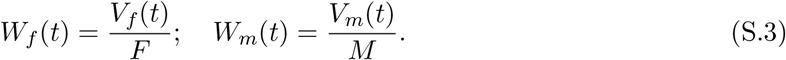

As a check, we may integrate these fitness functions across the female- and male-producing temperature ranges respectively; this returns the same value, confirming that the total male and female reproductive outputs are equal:

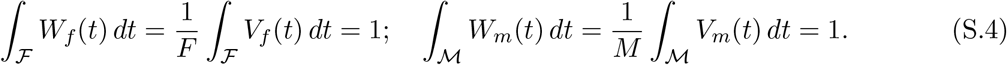

### ESS conditions

For the ESD system to be stable, its thresholds Θ^∗^ and corresponding sex-determining rule (ℱ^∗^, ℳ^∗^) must satisfy certain requirements set by *W*_*f*_ (*t*) and *W*_*m*_(*t*). Since *W*_*f*_ (*t*) and *W*_*m*_(*t*) are themselves determined by the parameters of the system, we write them in the context of the candidate ESS system Θ^∗^ as *W*_*m*_(*t* | Θ^∗^) and *W*_*f*_ (*t* | Θ^∗^), and define *D*(*t*) = *W*_*m*_(*t* | Θ^∗^)−*W*_*f*_ (*t* | Θ^∗^), the difference in the expected fitnesses of a male and a female offspring incubated at *t*.

**Condition 1**. First, for each threshold *θ*^∗^ ∈ Θ^∗^, the expected fitnesses of a male and a female offspring incubated at temperature *θ* must be the same: *D*(*θ*^∗^) = 0. Conversely, for each temperature *t* at which male and female fitnesses are equal (*D*(*t*) = 0), there must be a threshold *θ*^∗^ = *t* in the system. That is, the male and female fitness functions must cross at each threshold in the system, and there must be a threshold at each such crossing.

**Condition 2**. Second, for each threshold *θ*^∗^, if *D*^′^(*θ*^∗^) *>* 0, then females must be produced in the interval below *θ*^∗^ and males must be produced in the interval above *θ*^∗^. If, on the other hand, *D*^′^(*θ*^∗^) *<* 0, then males must be produced in the interval below *θ*^∗^ and females must be produced in the interval above *θ*^∗^.

Jointly, these conditions ensure that (and are required so that), at any temperature *t*, the higher-fitness sex at that temperature is produced; that is, *W*_*m*_(*t* | Θ^∗^) ≥ *W*_*f*_ (*t* | Θ^∗^) for all *t* ∈ ℳ^∗^, and *W*_*f*_ (*t* | Θ^∗^) ≥ *W*_*m*_(*t* | Θ^∗^) for all *t* ∈ ℱ^∗^ (see Main Text Figs. 2, 4).

## S2 Fitness cost of a threshold-biasing allele

In a resident system with ESS thresholds Θ^∗^, we consider one of the thresholds, *θ*^∗^. We assume, without loss of generality, that *D*^′^(*θ*^∗^) *>* 0 in the resident system, so that females are produced in the interval below *θ*^∗^ and males in the interval above *θ*^∗^.

Suppose that a rare allele increases the threshold of its bearers by an amount *δ*, from *θ*^∗^ to *θ*^∗^ +*δ*. We assume that, if the system is a multi-threshold one, this shift does not push the threshold above any other threshold. Therefore, across the temperature range (*θ*^∗^, *θ*^∗^ + *δ*], males have higher fitness than females (since *D*^′^(*θ*^∗^) *>* 0), but individuals carrying the allele develop as female instead of male, suffering an average fitness reduction

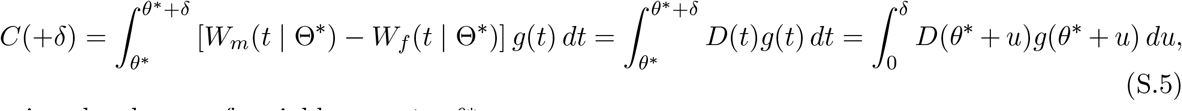

using the change of variables *u* = *t* − *θ*^∗^.

If the threshold shift *δ* is small, we may approximate the integrand by its first-order Taylor expansion

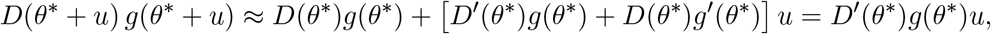

where we have used the ESS condition *D*(*θ*^∗^) = 0. So

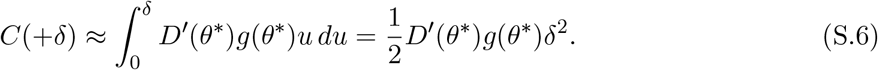

By a similar calculation, an allele that decreases its bearers’ thresholds from *θ*^∗^ to *θ*^∗^ − *δ* would suffer an average fitness cost

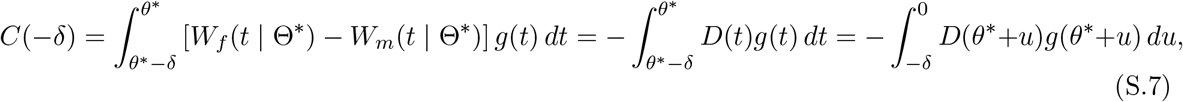

which, if the threshold shift is small, is again approximately

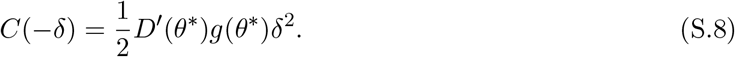

So, for small threshold shifts, the cost grows quadratically with the magnitude of the shift.

## S3 Invasion of a threshold-biasing, sexually antagonistic haplotype

We now consider a haplotype *H* containing a threshold-biasing allele *A* and a viability-affecting allele *B*. Label the resident alleles at the two loci *a* and *b*, and the resident *a*-*b* haplotype *h*.

*B*’s effects on viability are assumed to be multiplicative: compared to the viability *V*_*m*_(*t*) of a resident *bb* homozygote incubated at temperature *t*, a *Bb* heterozygote has viability (1 + *s*_*m*_)*V*_*m*_(*t*) while a *BB* homozygote has viability (1+*s*_*m*_)^2^*V*_*m*_(*t*); similarly, the viabilities of females of genotype *bb, Bb*, and *BB* incubated at temperature *t* are *V*_*f*_ (*t*), (1 + *s*_*f*_)*V*_*f*_ (*t*), and (1 + *s*_*f*_)^2^*V*_*f*_ (*t*) respectively. We restrict *s*_*m*_ and *s*_*f*_ to the interval [−1, 1]. We are particularly interested in the case where the viability-affecting allele is sexually antagonistic (*s*_*m*_ *<* 0, *s*_*f*_ *>* 0 or *s*_*m*_ *<* 0, *s*_*f*_ *>* 0), but we also allow it to have concordant effects on male and female viability.

*A* affects one threshold additively, increasing its value by *δ* in *Aa* heterozygotes and 2*δ* in *AA* homozygotes. Thus, when the haplotype is rare, and supposing the relevant threshold to be at its ESS value *θ*^∗^ in the resident population, the thresholds of *Aa* and *AA* individuals are *θ*^∗^ + *δ* and *θ*^∗^ + 2*δ* respectively. We assume, without loss of generality, that *D*^′^(*θ*^∗^) *>* 0 in the resident system, so that males are produced in the interval above the threshold and females in the interval below. Thus, the threshold-affecting allele *A* is present more often in females than the wild-type *a*.

We assume no recombination between the two loci, so that the haplotype *H* behaves as a single threshold-biasing, viability-affecting allele. We are interested in the conditions under which it can invade the resident population.

When *H* is rare, it is present only in *Hh* heterozygotes, with threshold *θ*^∗^ + *δ* and viabilities that differ from the residents (*hh*) by factors 1 + *s*_*m*_ and 1 + *s*_*f*_ . The resident ESS thresholds Θ^∗^ determine the population sex ratio and the the reproductive-value weights

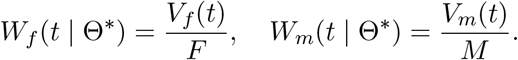

The mutant haplotype invades if the expected fitness of *Hh* heterozygotes in the resident background is greater than that of *hh* homozygotes. The expected fitness of the resident *hh* homozygotes is

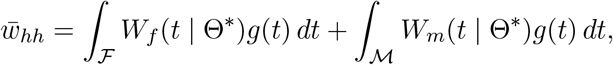

while the expected fitness of *Hh* heterozygotes is

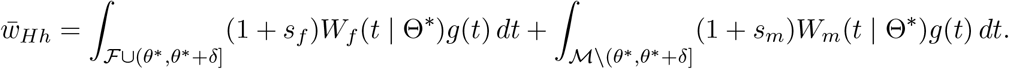

The difference is

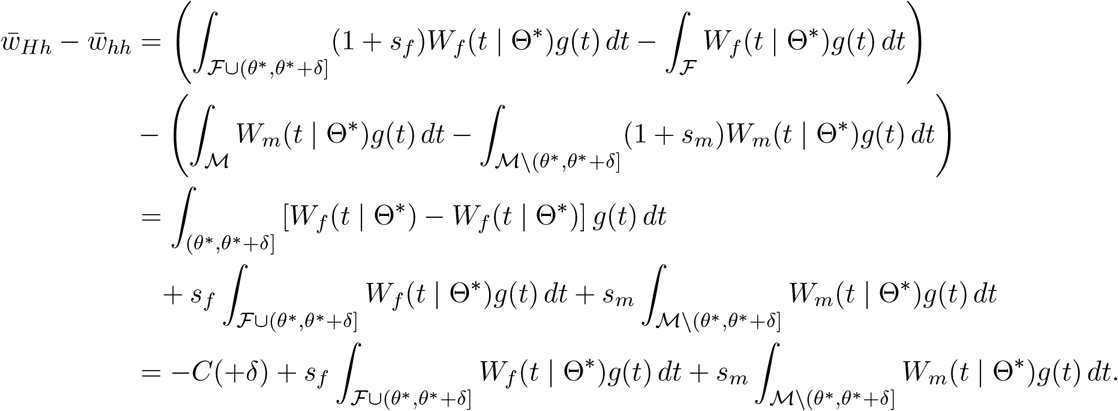

The *H* haplotype invades if 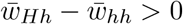; i.e., if

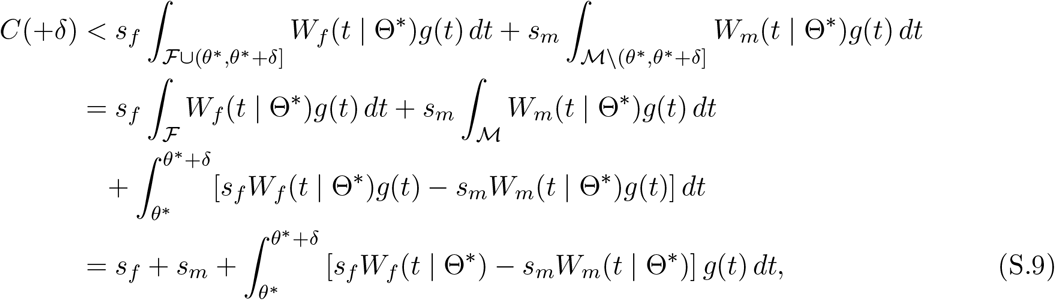

using Eq. (S.4).

If the threshold shift *δ* is small and the viability effects *s*_*m*_ and *s*_*f*_ are also small, specifically O(*δ*^2^), then the final integral in (S.9) is negligible, and we can use our earlier approximation for the fitness cost of a threshold shifted from its ESS value (Eq. S.6) to approximate the invasion condition (S.9) by

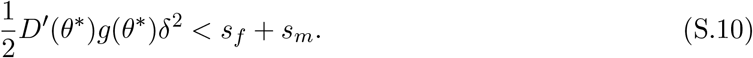

If, instead, the *H* haplotype decreases the threshold by an amount *δ*, and is therefore male-biasing, the expected fitness of *Hh* heterozygotes is

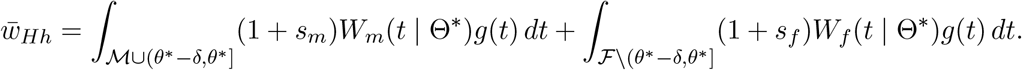

The difference is

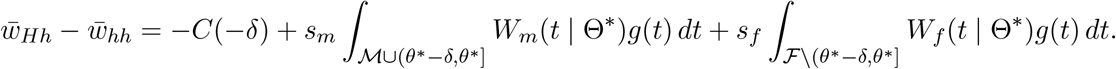

Therefore, the *H* haplotype invades if

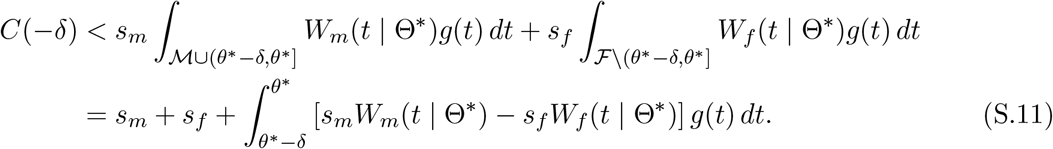

## S4 Conditions for polymorphism of a threshold-biasing, sexually antagonistic haplotype

If the haplotype *H* invades and spreads, the sex ratio of the population would, all else equal, become progressively more biased towards the sex overproduced by *H*. This would generate selection against *H* and check its spread. If, however, there was genetic variation for the threshold phenotype in the resident system, then the ‘background’ threshold value (i.e., excluding the contribution of *H*) would adapt in response to *H*’s spread. This leads to frequency-dependent selection on *H*, and so invasion from rarity does not necessarily guarantee fixation. Here, we derive the conditions under which the invading haplotype will spread to an intermediate frequency.

The logic is straightforward. We have derived above the condition for invasion of the haplotype *H* in a population initially fixed for *h*. Assuming that condition to be satisfied, we then imagine a population in which *H* is fixed, and ask whether the *h* haplotype can invade this population. If so, we conclude that the conditions support eventual polymorphism, since *H* invades but cannot fix.

### ESS thresholds in a population fixed for the threshold-biasing, sexually antagonistic haplotype

To calculate the conditions for invasion of *h* in a resident system fixed for *H*, we need to know what the ESS thresholds would be in the population fixed for *H*. Fortunately, these turn out to be the same as those in a population fixed for *h*. To see why, let the ESS sets of male- and female-determining temperatures in a population fixed for *h* be ℳ and ℱ. The thresholds Θ^∗^ that delimit these regions satisfy the ESS conditions set out in Section S1 above; in particular, for each such threshold *θ*^∗^, *W*_*m*_(*θ*^∗^ | Θ^∗^) = *W*_*f*_ (*θ*^∗^ | Θ^∗^), where the fitness functions *W*_*m*_ and *W*_*f*_ are implicitly determined by ℳ, ℱ, and the viability functions *V*_*m*_(*t*) and *V*_*f*_ (*t*) that pertain in the population fixed for *h*.

In a population fixed for *H, all* males have their viabilities multiplied by the same factor (1 + *s*_*m*_)^2^, and *all* females have their viabilities multiplied by the same factor (1 + *s*_*f*_)^2^, relative to what they would be in a population fixed for *h*. Assume the sets of male and female producing temperatures to be ℳ and ℱ, which we know by assumption to be ESS in a population fixed for *h*. In the population fixed for *H*, and given ℳ and ℱ, for each sex, both the viability function and the reproductive weight defined in Eq. (S.2) are multiplied by a constant factor ((1 + *s*_*m*_)^2^ for males, (1 + *s*_*f*_)^2^ for females), and so the fitness function defined by Eq. (S.3) is unchanged. Since the fitness functions given ℳ and ℱ are the same for a population fixed for *h* and a population fixed for *H*, and since ℳ and ℱ are ESS under these fitness functions (as we assumed for the population fixed for *h*), ℳ and ℱ (and the thresholds that define them) are ESS in a population fixed for *H* as well. Given genetic variation in the background threshold, it will adapt in response to the fixed *H* haplotype, precisely countering the effect of *H* on the threshold to restore the ESS values Θ^∗^.

### Conditions for reverse invasion of the wild-type haplotype in a population fixed for the threshold-biasing, sexually antagonistic haplotype

We focus on the case where *H* is threshold increasing, and we further assume that females are produced in the interval below the threshold that *H* affects, and males in the interval above.

Assume that *H* is fixed, and that the male- and female-determining temperatures are ℳ and ℱ, which we know to be ESS. We focus on the threshold that *h* and *H* affect, *θ*^∗^, which is an ESS threshold both when *h* is fixed and when *H* is fixed, following the logic above. In the population initially fixed for *H*, a rare haplotype *h* is present only in heterozygotes, and decreases their threshold from *θ*^∗^ to *θ*^∗^ − *δ*. Since *H* changes the viabilities of its male and female bearers multiplicatively by factors 1 + *s*_*m*_ and 1 + *s*_*f*_, relative to *h*, we can reciprocally think of *h* as changing the viabilities of its male and female bearers multiplicatively by factors 1*/*(1 + *s*_*m*_) and 1*/*(1 + *s*_*f*_), relative to *H*. That is, we can think of the male and female viability coefficients of *h* as

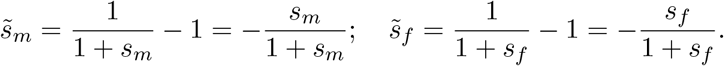

Since the ESS thresholds in the resident *H*-population are the same as they were when *h* was fixed, the invasion condition for *h* in the *H*-population can be calculated identically to the invasion condition for *H* in the *h*-population above (Eqs. S.9 and S.11). Therefore, the threshold-decreasing haplotype *h* can invade if

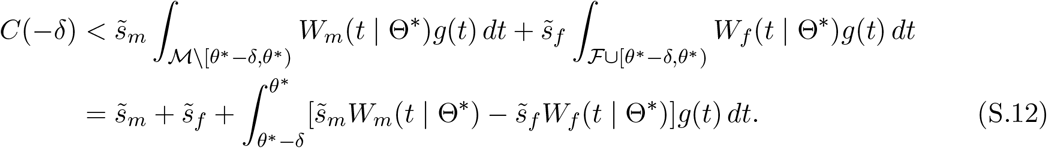

where

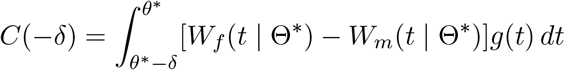

If the threshold shift *δ* is small and the viability effects 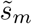 and 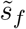 are O(*δ*^2^), then, recalling that for small *δ* we have 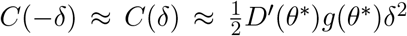, the invasion condition (S.9) can be approximated by

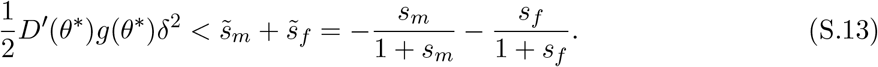

If *H* were instead threshold decreasing, then *h* is relatively threshold increasing, and so the condition for its reverse invasion is

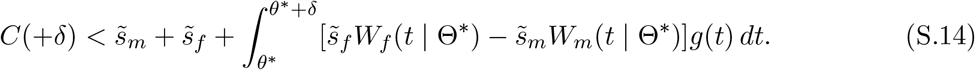

When *δ, s*_*m*_, and *s*_*f*_ are small, this again can be approximated by Eq. (S.13).

### Conditions for polymorphism

If the conditions (S.9) and (S.12) [or (S.11) and (S.14)] both hold, then the population will eventually be polymorphic for *H* and *h*. If condition (S.9) holds but condition (S.12) does not [or (S.11) does and (S.14) does not], then we expect *H* to invade and spread to fixation. We have not formally proven this latter statement, and will not attempt to do so, but our simulations suggest that it is true (Figs. 3, 4, S3).

## S5 The equilibrium frequency of the threshold-biasing, sexually antagonistic allele

When the sexually antagonistic, threshold-biasing haplotype *H* invades but cannot fix, what is the equilibrium frequency it attains (if any)? To characterize this, we need to make assumptions about the response of other loci that affect the threshold to the spread of *H*. As *H* spreads, all else equal, it would bias the sex ratio of the population towards the sex it overproduces; this creates selection for adjustment of the ‘background’ threshold value to produce more of the other sex. Whether or not, and how fast, the background threshold responds to this selection depends on its genetic basis.

In particular, we assume that the background threshold is a highly polygenic quantitative trait, as evidence indicates it often is. It will therefore always present substantial genetic variance, and will therefore always adapt to changes in the frequency of the *H* haplotype. How rapidly it adapts in response to *H*’s spread will depend on (i) the genetic variance for the background threshold, which we assume to be a fixed value, and (ii) the selection gradient acting on the background threshold for a given frequency of *H*, which we calculate below. At equilibrium, the selection gradients on the haplotype’s frequency and the background threshold must be zero.

As before, we assume that males are produced in the interval above the relevant threshold, with females produced in the interval below, and that the haplotype *H* increases the threshold, by *δ* in heterozygotes and 2*δ* in homozygotes; the case where it decreases the threshold can be treated similarly. Given a value of the ‘background’ threshold *θ*_bg_, the threshold phenotypes of the three genotypes *hh, Hh*, and *HH* are *θ*_bg_, *θ*_bg_ + *δ*, and *θ*_bg_ + 2*δ* respectively. If the frequency of *H* among adult males and females in the previous generation was *p*_*m*_ and *p*_*f*_ respectively, and mating is random, then among zygotes in this generation, the three genotypes are found in frequencies *p*_*hh*_ = (1 − *p*_*m*_)(1 − *p*_*f*_), *p*_*Hh*_ = *p*_*m*_(1 − *p*_*f*_) + (1 − *p*_*m*_)*p*_*f*_, and *p*_*HH*_ = *p*_*m*_*p*_*f*_ (note that these frequencies will not necessarily be in Hardy–Weinberg proportions, since *p*_*m*_ and *p*_*f*_ can differ). As before, the viabilities of the three genotypes *hh, Hh*, and *HH* are multiplied by 1, 1 + *s*_*m*_, and (1 + *s*_*m*_)^2^ in males and 1, 1 + *s*_*f*_, and (1 + *s*_*f*_)^2^ in females.

From these values, the numbers of males and females that survive viability selection are proportional to

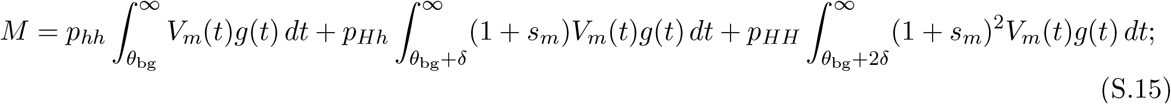

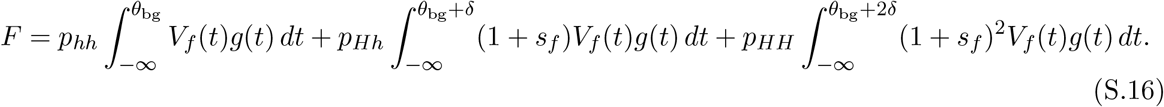

*M* and *F* determine the reproductive value of a surviving male (∝ 1*/M*) and a surviving female (∝ 1*/F*).

Consider a rare background threshold phenotype 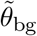 that differs from the resident value *θ*_bg_. The average fitness of this alternative background threshold in a population in which almost all individuals have *θ*_bg_ as their background threshold is

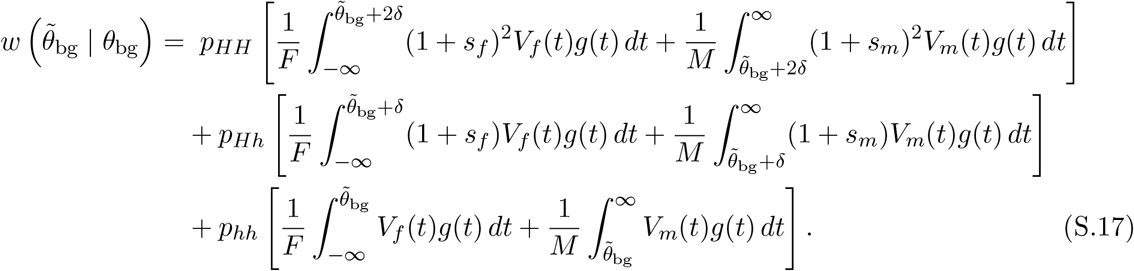

Taking the derivative (using the first fundamental theorem of calculus),

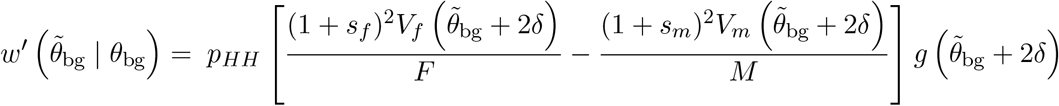

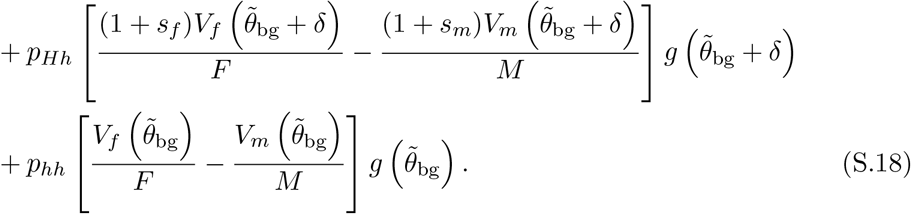

The selection gradient at *θ*_bg_ is therefore

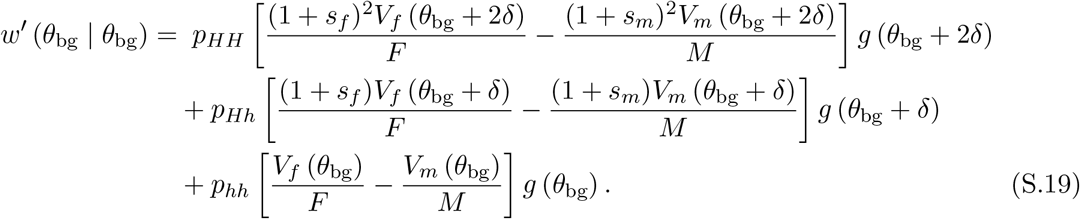

Note that the right-hand side of Eq. (S.19) is the average of the differences between female and male fitness of the three genotypes *HH, Hh*, and *hh* at their respective threshold temperatures, weighted by the density of the environmental distribution *g* at these threshold temperatures. The condition for (partial) equilibrium of the background threshold, given the genotype frequencies of *H* haplotype, is that the selection gradient on *θ*_bg_ be zero. This condition, that the weighted average fitnesses of males and females across the three thresholds must be equal, is analogous to the ESS condition for a threshold, Condition 1 in Section S1 above, and indeed, if *p*_*HH*_ = 0 or *p*_*hh*_ = 1, setting Eq. (S.19) to zero would recover this ESS condition.

The general (as opposed to partial) equilibrium is the set of frequencies 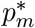 and 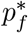 and the partial equilibrium value 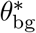 associated with those frequencies such that, assuming all individuals to have background threshold value 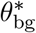, the frequencies 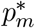 and 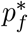 (measured in adults from the previous generation) remain constant from one generation to the next. We will not attempt to characterize this general equilibrium analytically, and instead describe here a numerical method to calculate the equilibrium. The method essentially simulates evolution of the system as the *H* haplotype invades (or goes extinct), assuming the background threshold to be a quantitative trait with a certain genetic variance *V*_*G*_; we take the values that the simulation converges to as the equilibrium values of the system.

Given a starting set of frequencies *p*_*m*_ and *p*_*f*_ (= 0.01 in the simulations we have run), and an initial value of the background threshold *θ*_bg_ (equal to its ESS value in our simulations), we calculate (i) the frequencies of the three genotypes among zygotes (*p*_*HH*_ = *p*_*m*_*p*_*f*_, etc.), (ii) the frequency of *H* among adult males and females this generation (based on the viabilities of the three genotypes and the environments in which they result in male or female development), and (iii) the selection gradient *S* on the background threshold, given by Eq. (S.19). We then move to the next generation, treating the adult frequencies from step (ii) as the new values 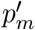 and 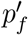, and with a new value of the background threshold 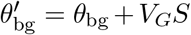 (in our simulations, we set *V*_*G*_ = 0.02). We iterate this procedure until convergence, and treat the values 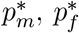, and 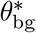 that the simulation converges to for given set of parameters as the equilibrium values for that parameter set.

Given the equilibrium frequencies 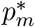 and 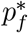 of the haplotype among adults and the number of males and females in the adult population, we calculate the average frequency of the haplotype among adults. This is the quantity plotted in Figs. 3, 4, and S3.

## S6 Parameters used in the figures

The details of the viability functions used in the Main Text figures are as as follows:

- **Single-threshold case, linear viability**. Female viability increases linearly with temperature, *V*_*f*_ (*t*) = *t/*2 + 4*/*9, while male viability increases more steeply, *V*_*m*_(*t*) = (12*/*9)*t*. These functions yield an ESS threshold value *θ*^∗^ = 0.52 and equilibrium sex ratio (fraction males) *m*^∗^ = 0.44.
- **Single-threshold case, logistic male viability**. Female viability remains *V*_*f*_ (*t*) = *t/*2+4*/*9, while male viability follows *V*_*m*_(*t*) = *λ*{1 + exp[−*k*(*t* − 0.5)]}^−1^ with *k* = 20 and *λ* = 12*/*9, scaled to match the linear case at *t* = 1. This yields *θ\** = 0.53 and *m\** = 0.41.
- **Two-threshold case, opposing Gaussian fitness functions**. Male viability follows a Gaussian hump and female viability a Gaussian trough centered on 1*/*2 and with parameters (base, amp, *σ*) = (0.10, 0.90, 0.08) for males and (0.20, 0.80, 0.10) for females. These functions yield 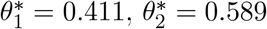, and *m*^∗^ = 0.522.

Unless otherwise stated, the environmental distribution is normal with *µ* = 0.5 and *σ* = 0.125, but truncated to the range [0, 1] and rescaled so that it integrates to 1. In the figures, the threshold-biasing allele reduces the threshold and is therefore male-biasing in all cases. Analyses were conducted in R using RStudio (v2024.04). Code used to generate the results and figures is available on GitHub: github.com/Pavitra451/Charnov_Bull_sexual_antagonism.

## Supplementary Figures

**Figure S1.**
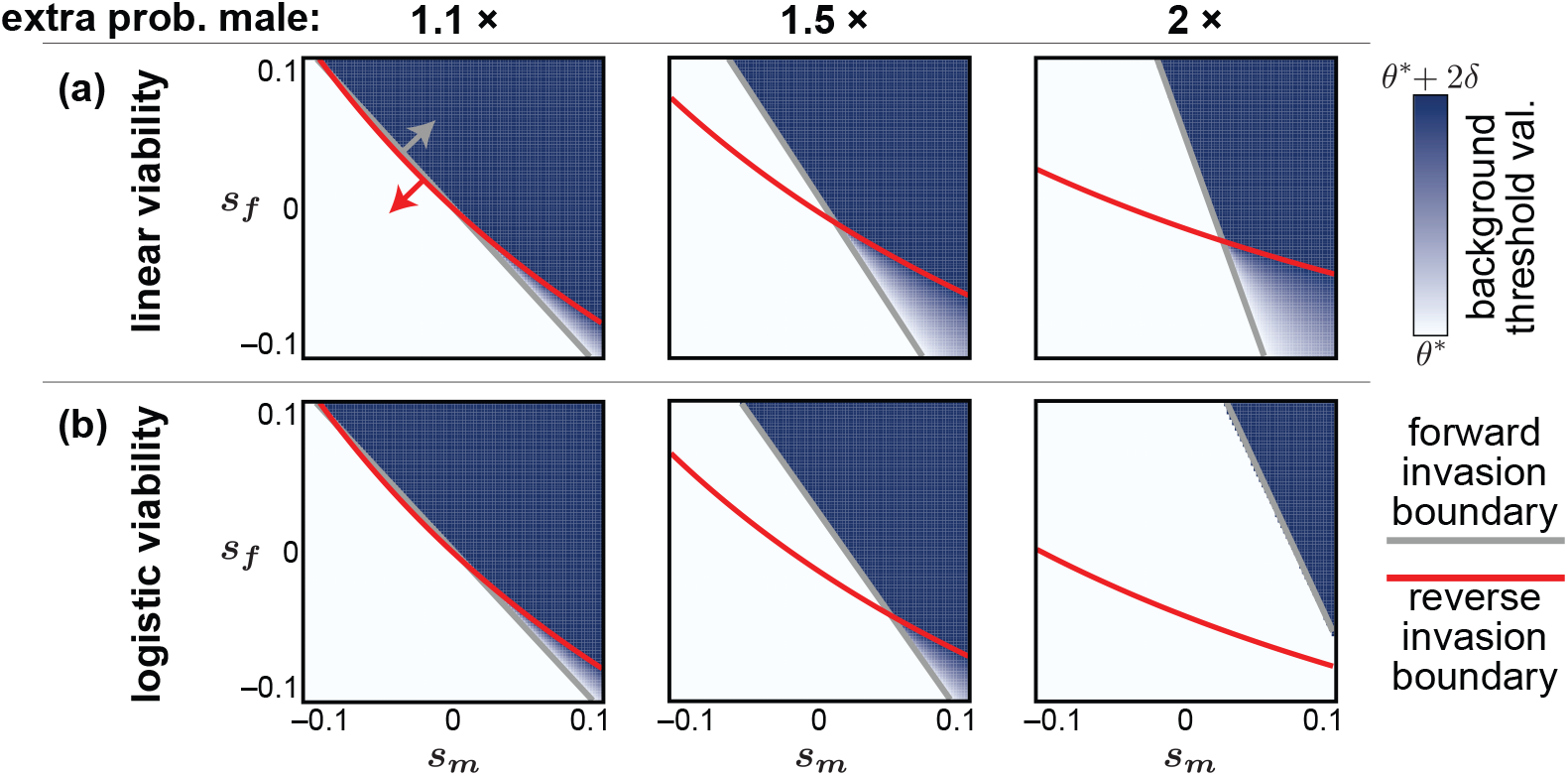
The background threshold phenotype evolves to accommodate the spread of threshold-biasing alleles. As in Fig. 3, but here the value of the background threshold phenotype is shown rather than the equilibrium frequency of the threshold-decreasing haplotype *H*. In regions where *H* cannot invade and is lost, the background threshold remains at its original ESS value. In regions where the allele fixes, the background threshold evolves to the original ESS value plus 2*δ*, to exactly counter the effect of *H* in homozygotes, −2*δ*. The overall threshold in regions of loss and fixation of *H* therefore returns to the original ESS. In contrast, in regions where the allele maintains an equilibrium frequency between 0 and 1, the overall threshold phenotype deviates from the original ESS.

**Figure S2.**
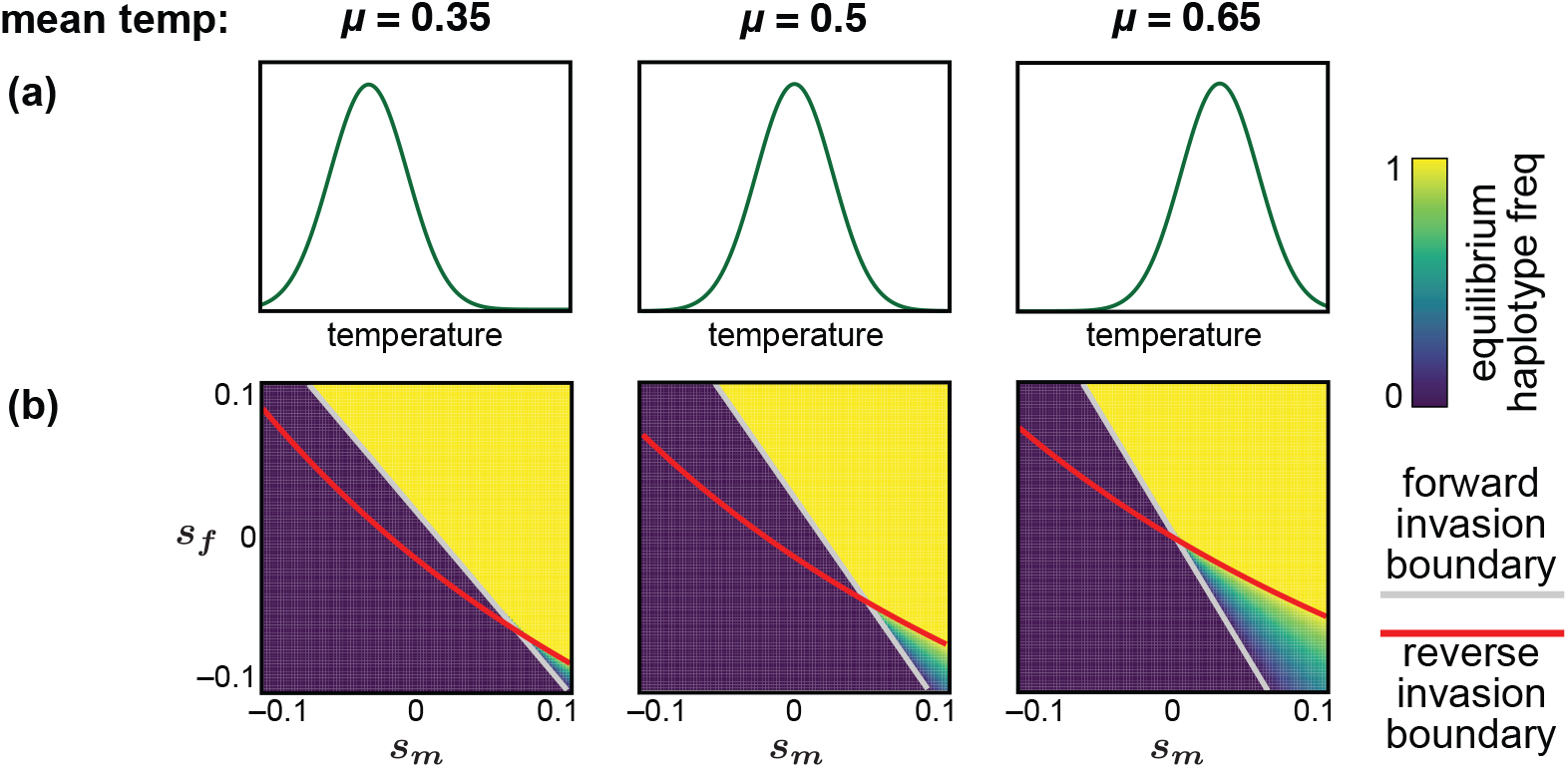
The distribution of environments affects the Charnov–Bull costs associated with a threshold-biasing haplotype, and therefore also the potential for invasion and stable polymorphism of the haplotype. As in Fig. 3, but with the environmental distribution centered at *µ* = 0.35 (left), *µ* = 0.5 (center; identical to Fig. 3), and *µ* = 0.65 (right). When *µ* is lowered, more weight of the environmental distribution falls within the region affected by the threshold shift, increasing Charnov–Bull costs and causing the parameter space permitting stable polymorphism to shrink. In contrast, when less weight is placed in that region, Charnov–Bull costs are reduced and the parameter space supporting polymorphism expands.

**Figure S3.**
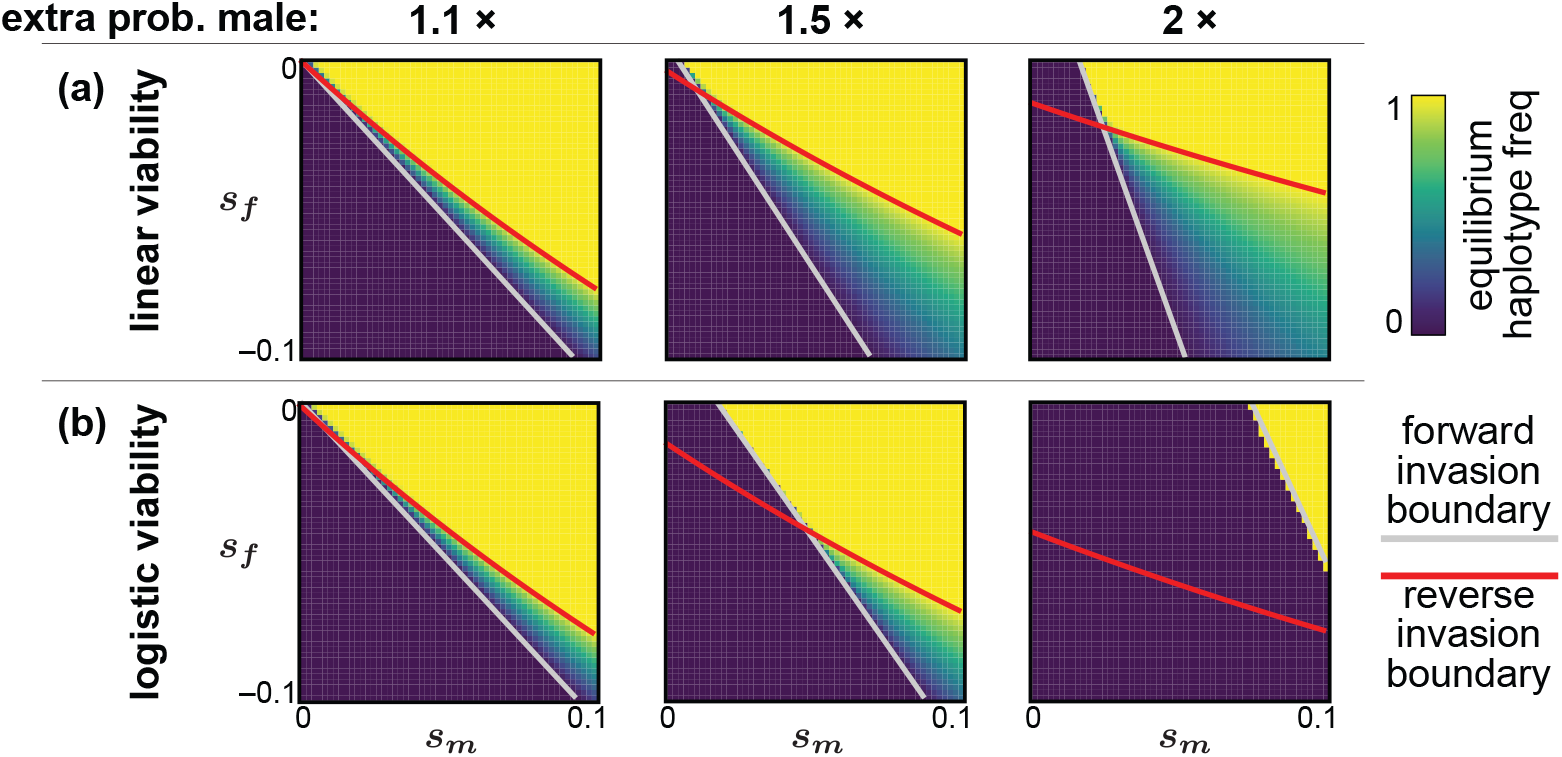
As in Fig. 3, but only the sexually antagonistic portion of parameter space is shown. Because the haplotype’s threshold effect is male-biasing here, invasion is most likely when it is male beneficial, female costly. Accordingly, we depict only the region where *s*_*f*_ *<* 0 (ranging from 0 to −0.1) and *s*_*m*_ *>* 0 (ranging from 0 to 0.1).

## Notes

### Competing Interest Statement

The authors have declared no competing interest.

### Summary of Updates

Axes on the right panels in Fig. 2b and c were inadvertently reversed in previous version.

